# APOBEC mutagenesis inhibits breast cancer growth through induction of a T cell-mediated antitumor immune response

**DOI:** 10.1101/2021.02.13.431068

**Authors:** Ashley V. DiMarco, Xiaodi Qin, Sarah Van Alsten, Brock McKinney, Nina Marie G. Garcia, Jeremy Force, Brent A. Hanks, Melissa A. Troester, Kouros Owzar, Jichun Xie, James V. Alvarez

**Affiliations:** Department of Pharmacology and Cancer Biology, Duke University School of Medicine; Department of Biostatistics and Bioinformatics, Duke University School of Medicine; Department of Epidemiology, Gillings School of Global Public Health, University of North Carolina at Chapel Hill; Division of Medical Oncology, Department of Medicine, Duke Cancer Institute

**Keywords:** APOBEC, mutational signatures, APOBEC3B, breast cancer, immunotherapy, checkpoint blockade

## Abstract

The APOBEC family of cytidine deaminases is one of the most common endogenous sources of mutations in human cancer. Genomic studies of tumors have found that APOBEC mutational signatures are particularly enriched in the HER2 subtype of breast cancer and have been associated with immunotherapy response in diverse cancer types. However, the direct consequences of APOBEC mutagenesis on the tumor immune microenvironment have not been thoroughly investigated. To address this, we developed syngeneic murine mammary tumor models with inducible expression of APOBEC3B. We found that APOBEC activity induces an antitumor adaptive immune response and CD4^+^ T cell-mediated tumor growth inhibition. While polyclonal APOBEC tumors had a moderate growth defect, clonal APOBEC tumors were almost completely rejected by the immune system, suggesting that APOBEC-mediated genetic heterogeneity limits the antitumor adaptive immune response. Consistent with the observed immune infiltration in APOBEC tumors, APOBEC activity sensitized HER2-driven breast tumors to checkpoint inhibition. In human breast cancers, the relationship between APOBEC mutagenesis and immunogenicity varied by breast cancer subtype and the frequency of subclonal mutations. This work provides a mechanistic basis for the sensitivity of APOBEC tumors to checkpoint inhibitors and suggests a rationale for using APOBEC mutational signatures as a biomarker predicting immunotherapy response in HER2-positive breast cancers.

**SIGNIFICANCE:** APOBEC mutational signatures are observed in many cancers, yet the consequences of these mutations on the tumor immune microenvironment are not well understood. Using a novel mouse model, we show that APOBEC activity sensitizes HER2-driven mammary tumors to checkpoint inhibition and could inform immunotherapy treatment strategies for HER2-positive breast cancer patients.

## INTRODUCTION

More than 50 distinct mutational signatures have been identified in cancer genomes (1–3). These signatures are thought to reflect transient or ongoing exogenous and endogenous mutational processes that occur over the lifetime of normal cells and during tumor development. Single-base substitution (SBS) signature 2 is characterized by C-to-T transitions within the trinucleotide motif of TCW (where W represents adenine or thymine), and SBS signature 13 is defined by C-to-G transversions within the same TCW motif. Both signatures are attributed to the APOBEC (apolipoprotein B mRNA editing enzyme, catalytic polypeptide-like) family of cytidine deaminases (1, 2). APOBEC enzymes catalyze the deamination of cytosine to uracil on single-stranded DNA which, following repair, manifests predominantly as C-to-T and C-to-G point mutations. The majority of these substitutions are distributed stochastically throughout the somatic genome; however, some are localized in multi-kilobase long, strand-coordinated clusters referred to as ‘kataegis’ (1,4–6). While APOBEC enzymes have evolutionarily conserved activity in the generation of antibody diversification and restriction of viruses and endogenous retrotransposons, their off-target mutagenic activity on the host somatic genome drives cancer genome instability (reviewed by (7, 8)). APOBEC-mediated mutational signatures have been detected in at least 22 different tumor types and are particularly enriched in bladder, head and neck, cervical, and breast cancer (9, 10). Importantly, nearly half of breast cancers exhibit kataegis hypermutation clusters (11). Among breast cancer subtypes, human epidermal growth factor receptor 2-positive (HER2^+^) breast tumors are reported to have the highest median levels of APOBEC signature enrichment (2,9,12).

Somatic mutations in cancer can give rise to unique mutant peptides that serve as immune-reactive neoantigens, allowing cytotoxic T cells to target tumor cells for elimination (13, 14). Thus, recent work has focused on understanding the role of ongoing mutational processes in contributing to tumor immunogenicity and response to immunotherapies (15–17). Despite the prevalence of APOBEC mutational signatures in breast cancer, these tumors are traditionally thought to be poorly immunogenic or “cold”. Breast tumors generally have a modest tumor mutation burden (TMB) (2) and low tumor-infiltrating lymphocytes relative to more immunogenic cancers that exhibit robust immune infiltration and are sensitive to checkpoint inhibitors (reviewed by (18–20)). However, the initial trials of anti-PD-1 and anti-PD-L1 monotherapy in triple-negative breast cancer (TNBC) showed promising objective response rates of up to 19% (21, 22). The combination of anti-PD-L1 and nab-paclitaxel had an objective response rate of 39% and prolonged overall survival, leading to its FDA approval for advanced/metastatic PD-L1^+^ TNBC in 2019, the first approval of immunotherapy for breast cancer (23). However, checkpoint inhibitor clinical trials have been less successful for HER2^+^ breast cancer patients. In an initial trial for anti-PD-L1 monotherapy in metastatic breast cancer, there were no objective responses in the HER2^+^ subtype (24). When trastuzumab was combined with anti-PD-1 for HER2^+^ patients, responses ranged from 0%-15.2% and were highly dependent on PD-L1 status (25). However, APOBEC mutational signatures have yet to be investigated as a specific class of hypermutation that transforms an immunologically “cold” HER2^+^ breast tumor “hot”, rendering the tumor responsive to checkpoint inhibition.

Recent work on how mutational signatures impact tumor immunity has revealed several pieces of evidence potentially implicating APOBEC mutagenesis in immunotherapy response. In pan-cancer analyses from The Cancer Genome Atlas (TCGA), the kataegis-like APOBEC mutational signature was significantly correlated with PD-L1 expression and neopeptide hydrophobicity (26, 27). Further, APOBEC signatures were associated with a greater likelihood of response to immune checkpoint inhibition in non-small cell lung cancer (NSCLC) (28), head and neck cancer, bladder cancer (29), and in a small cohort of breast cancer patients (30). In a recent study using mouse models of TNBC, overexpression of the murine APOBEC3 ortholog sensitized tumors to checkpoint inhibitors (31). Additionally, overexpression of human APOBEC3B in a vaccine setting sensitized mouse melanomas to checkpoint inhibition (32). However, the direct consequences of APOBEC mutagenesis on the tumor immune microenvironment and tumor growth in the absence of checkpoint inhibitors have not been thoroughly explored. A mechanistic understanding of how APOBEC mutagenesis alters the tumor immune microenvironment would inform the use of immune therapies for human tumors with APOBEC mutational signatures. Furthermore, despite the high enrichment of APOBEC signatures in HER2^+^ breast cancer, no studies to our knowledge have investigated a role for APOBEC mutagenesis in conferring clinical benefit to checkpoint blockade in HER2^+^ breast cancer.

To address these questions, we developed a syngeneic, immunocompetent murine HER2-driven mammary tumor model with APOBEC activity. Using this model, we examined the consequences of APOBEC activity and genetic heterogeneity on tumor growth, investigated tumor-immune system interactions in APOBEC tumors, and assessed the therapeutic response of these tumors to checkpoint inhibitor therapy. Finally, we examined the relationship between APOBEC mutagenesis and adaptive immune response in human breast tumors.

## RESULTS

### Ectopic expression of A3B in murine mammary tumor cells is not lethal and induces cytidine deaminase activity

To induce APOBEC mutagenesis *in vivo* in an immunocompetent HER2-driven mammary tumor model, we utilized the SMF cell line, which is derived from a mammary tumor arising in the MMTV-Neu/Her2 mouse model on the FVB background (33). We engineered SMF cells to conditionally express the human APOBEC family member, APOBEC3B (A3B), and thereby acquire APOBEC mutational signatures during tumor progression. Along with APOBEC3A, A3B is one of the major contributors of APOBEC mutations in cancer genomes (3,10,34,35). Studies in yeast and mammalian cells have shown that expression of A3B is sufficient to induce a kataegis-like pattern, and preferentially induce mutations at the TCW trinucleotide context resembling SBS signatures 2 and 13 in cancer genomes (5,36,37), whereas the murine APOBEC3 ortholog localizes to the cytoplasm and has low catalytic activity (38). The SMF cell line was stably transduced with a lentivirus encoding reverse tetracycline-controlled transactivator (rtTA) and a lentivirus encoding rtTA-responsive human A3B (referred to as “SMF-A3B cells”). This system allows for titratable and reversible expression of A3B in tumor cells with the administration of doxycycline (dox) in the cell culture medium or in the drinking water of mice.

To characterize the A3B expression system *in vitro*, SMF-A3B cells were cultured with increasing concentrations of dox, with or without subsequent removal of dox from the medium. A3B mRNA and protein expression were dose-responsive and reversible (Fig. 1A, B). Additionally, A3B protein was constitutively localized to the nucleus in the presence of dox, demonstrating proper subcellular localization of this APOBEC family member (39)(Fig. 1C). In an *in vitro* cytidine deaminase activity assay, increasing concentrations of dox induced dose-responsive deaminase activity in SMF-A3B cells (Fig. 1D). A3B expression did not affect cell proliferation or survival, as measured by an ATP-based cell viability assay and colony formation assay (Fig. 1E-G). For subsequent experiments we used a dox concentration (1 μg/mL) that induced A3B expression levels and deaminase activity levels comparable to that of the APOBEC-high human HER2^+^ breast cancer cell line, BT474 (Fig. 1A, D). These data suggest SMF-A3B is a suitable system to induce A3B expression and cytidine deaminase activity in a syngeneic, orthoptic murine tumor model.

**Figure 1:**
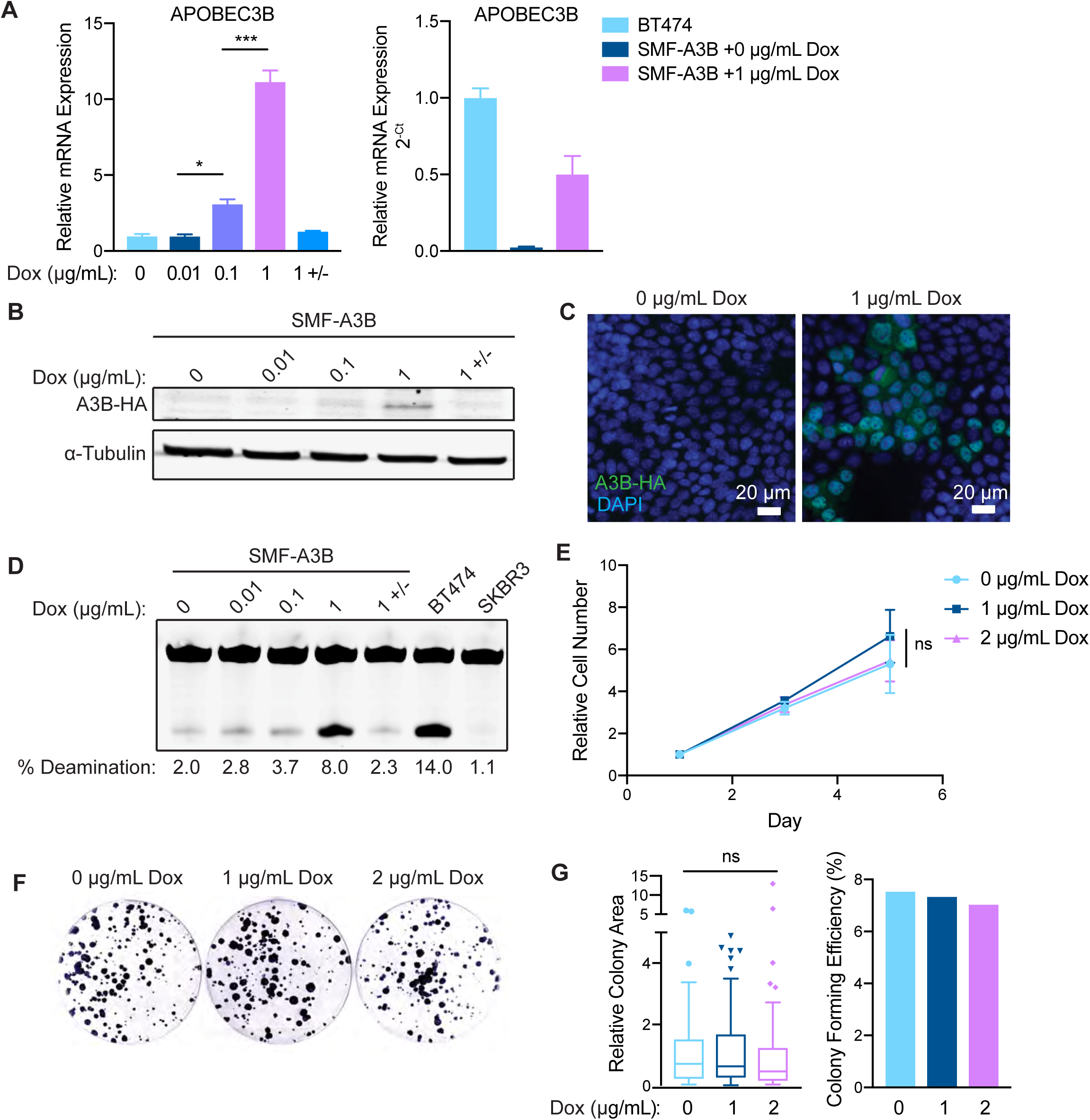
SMF-A3B cells express titratable and reversible APOBEC3B without loss in cell viability. **(A)** qRT-PCR of A3B gene expression in SMF-A3B cells treated with the indicated concentrations of dox for 2 days. 1 +/− indicates treatment with 1 µg/mL of dox for 2 days, then removal of dox for 2 days. Left: A3B expression relative to 0 µg/mL dox condition. Right: A3B expression relative to BT474 cells. Data are representative of 2 independent experiments. Results show 3 biological replicates and error bars depict mean ± SEM. Significance was determined using a one-way ANOVA and Tukey’s multiple comparisons test. **(B)** Western blot of HA-tagged A3B in SMF-A3B cells treated with dox as in (A). **(C)** Immunofluorescence staining for the HA epitope in SMF-A3B cells treated with dox as in (A), showing nuclear localization of HA-A3B. Blue channel is DAPI and green channel is HA. **(D)** In vitro cytidine deaminase activity assay of SMF-A3B cells treated with dox as in (A). The APOBEC-high human cell line BT474 and the A3B-null human cell line SKBR3 are shown as controls. **(E)** CellTiter-Glo assay showing growth curves of SMF-A3B cells treated with the indicated concentration of dox. Results are shown as mean ± SD of 3 replicates. Statistical significance was determined by two-way ANOVA. **(F)** Clonogenic assay of SMF-A3B cells cultured with dox for 2 weeks to measure long term survival. Colonies were stained with crystal violet. **(G)** Quantification of clonogenic assay in (F). Left: Boxplots depicting the relative colony area. Statistical significance was determined using a one-way ANOVA. Right: Colony forming efficiency in each condition. ns p > 0.05, * p < 0.05, ** p < 0.01, *** p < 0.001

### A3B expression does not affect tumor growth in immunodeficient mice and is predicted to induce APOBEC-mediated mutational signatures

We first tested whether expression of A3B affects tumor growth in the absence of an adaptive immune system. SMF-A3B cells were implanted in the mammary fat pad of immunodeficient NOD.Cg-*Prkdc^scid^ Il2rg^tm1Wjl^*/SzJ (NSG) mice, and mice received either normal or dox drinking water throughout the duration of tumor growth to express A3B. Control and A3B-expressing tumors grew at similar rates (Supplementary Fig. S1A), indicating that in the absence of a functional immune system, A3B expression does not affect tumor growth. This is consistent with the finding that A3B expression does not affect the growth or viability of SMF cells *in vitro*.

We next examined whether SMF-A3B tumors have evidence of APOBEC mutagenesis. Because APOBEC-induced mutations are randomly distributed throughout the genome, it is technically difficult to detect these mutations in a heterogeneous population of cancer cells (36, 40). Therefore, to measure the APOBEC mutational process in mouse tumors generated from the SMF-A3B cell line, we developed a gene expression-based classifier for prediction of APOBEC mutational signatures. The classifier was trained using sets of differentially expressed genes from RNA-seq data of APOBEC-high and APOBEC-low breast cancers from TCGA using 10-fold cross validation (as determined by APOBEC mutational signature enrichment score from whole-exome sequencing; see Methods). This analysis suggested that a 10-gene classifier was optimal for prediction. The genes selected for the classifier were *AXIN2*, *CCDC157*, *ICOS*, *NAGS*, *NXPH3*, *PRODH*, *PSD*, *SRRM3*, *STMN3*, and *TTC25*.

We next used this classifier to test whether SMF-A3B tumors in NSG mice have evidence of APOBEC mutagenesis. We performed RNA-seq on 6 control tumors and 6 tumors expressing A3B. When applied to this independent dataset, the classifier correctly identified 4 of 6 A3B-expressing tumors as well as 4 of 6 control tumors (66% sensitivity and 66% specificity, Supplementary Fig. S1B), for an overall accuracy in mouse tumors of 66%. This indicates that the gene expression-based classifier may be used to predict APOBEC mutational signatures in the genomes of human and murine tumors, and A3B-expressing tumors generated from SMF-A3B cells are likely to harbor genomic APOBEC-mediated mutations.

### APOBEC activity slows mammary tumor growth and triggers the infiltration of antitumor adaptive immune cells

Given the evidence that the APOBEC mutational signature is associated with both an immune response and sensitivity to immunotherapy in NSCLC, bladder, and head and neck cancer, we examined the effects of *in vivo* APOBEC activity on the tumor immune microenvironment. SMF-A3B cells were orthotopically implanted bilaterally in the mammary gland of syngeneic, immunocompetent wildtype FVB mice. One cohort of mice was administered dox in the drinking water to induce A3B expression and APOBEC activity in the tumor cells throughout tumor growth (“APOBEC tumors”), while the control cohort received normal drinking water (Fig. 2A). Interestingly, APOBEC tumors grew significantly slower than control tumors and had a smaller mass at endpoint (Fig. 2B). Immunofluorescence staining of APOBEC and control tumors for a marker of double-stranded DNA breaks, γH2AX, showed no activation of the DNA damage response *in vivo* (Supplementary Fig. S1C, D). Similarly, A3B expression for two weeks did not induce γH2AX or cleaved PARP in SMF-A3B cells *in vitro* (Supplementary Fig. S1E). Taken together with the finding that A3B expression does not affect cell growth *in vitro* (Fig. 1E-G) or tumor growth in immunodeficient NSG mice (Supplementary Fig. S1A), this suggests that the growth defect of APOBEC tumors was mediated by a tumor cell-extrinsic mechanism, specifically the immune response.

**Figure 2:**
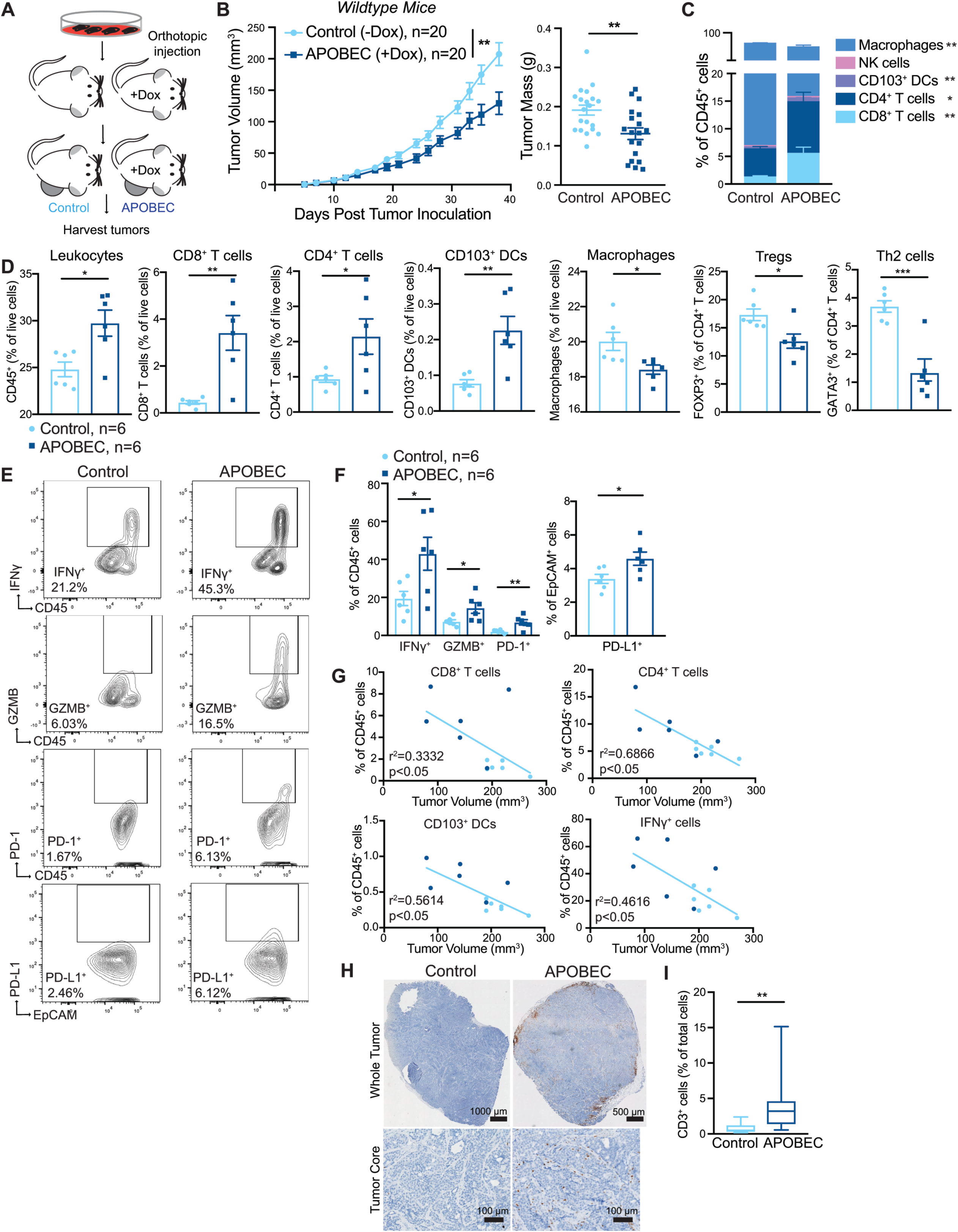
A3B expression slows tumor growth and triggers the infiltration of antitumor immune cells into the tumor core. **(A)** Schematic showing experimental design for tumor growth experiment. SMF-A3B cells were orthotopically implanted in the mammary gland of mice. The APOBEC cohort was administered dox in the drinking water and control cohort was administered normal drinking water until endpoint. **(B)** Left: tumor volume of control (n=20) and APOBEC tumors (n=20) in wildtype mice. Statistical significance was determined by two-way repeated-measures ANOVA. Right: tumor mass (g) of control and APOBEC tumors at endpoint. Statistical significance was determined by unpaired Student’s t-test. Error bars denote mean ± SEM. Data is representative of 2 independent experiments. **(C)** The frequency of immune cell types, expressed as a percentage of total CD45+ cells, in control (n=6) and APOBEC (n=6) tumors as determined by flow cytometry. **(D)** Flow cytometry quantification of immune cells in control (n=6) and APOBEC (n=6) tumors. Leukocytes, CD8+ T cells, CD4+ T cells, CD103+ dendritic cells (DCs), and macrophages are represented as the percentage of total live cells. T regulatory cells (Tregs) and type-2 T helper (Th2) cells are represented as the percentage of total CD4+ T cells. Statistical significance was determined by unpaired Student’s t-test. Error bars denote mean ± SEM. **(E-F)** Representative flow cytometry plots (E) and quantification (F) of staining for IFN, Granzyme B, PD-1 and PD-L1 in control (n=6) and APOBEC (n=6) tumors. Statistical significance was determined by unpaired Student’s t-test. Error bars denote mean ± SEM. **(G)** Pearson correlation between immune cell frequency and mean tumor volume (mm3) in control (light blue) and APOBEC (dark blue) tumors. Only significant correlations are shown, and the r squared values are indicated. **(H)** Immunohistochemistry (IHC) staining for the T cell marker CD3 in control and APOBEC tumors. The top image is a tiled scan of the whole tumor, and the bottom image is a representative region in the tumor core. **(I)** Quantification of CD3 staining for control (n=4) and APOBEC (n=4) tumors. Four fields of view were imaged for each tumor. Boxplots show the median percentage of CD3+ cells with minimum and maximum whiskers. Statistical significance was determined by unpaired Student’s t test with Welch’s correction. * p < 0.05, ** p < 0.01, *** p < 0.001

To gain insight into how A3B expression alters the tumor microenvironment (TME) of APOBEC tumors, six mice per cohort were randomly selected for immune profiling by flow cytometry (see Supplementary Fig. S2 for gating strategy and Supplementary Fig. S3 for representative FACS plots). APOBEC tumors showed a substantial infiltration of total leukocytes (CD45^+^EpCAM^−^) compared to control tumors (Fig. 2D). CD8^+^ T cells (CD45^+^CD3^+^CD8^+^), CD4^+^ T cells (CD45^+^CD3^+^CD4^+^), and CD103^+^ dendritic cells (DCs; CD45^+^CD11c^+^MHC-II^+^F4/80^−^ CD103^+^) were expanded in the APOBEC TME, as measured both as the percentage of CD45^+^ cells (Fig. 2C) and the percentage of total live cells (Fig. 2D). There was no change in the infiltration of natural killer (NK) cells (CD45^+^NK1.1^+^CD3^−^), although several subsets of cells that may have immunosuppressive potential were significantly reduced in the APOBEC tumors, including the fraction of T regulatory cells (Tregs; CD45^+^CD3^+^CD4^+^FOXP3^+^), type-2 T helper cells (Th2; CD45^+^CD3^+^CD4^+^GATA3^+^), and tumor-associated macrophages (CD45^+^F4/80^+^CD11c^low^) (Fig. 2D). Furthermore, APOBEC tumors were comprised of more immune cells producing the proinflammatory cytokine interferon-γ (IFNγ; CD45^+^IFNγ^+^), and cytotoxic granule granzyme B (GZMB; CD45^+^GzmB^+^) (Fig. 2E, F). Finally, PD-1^+^ immune cells (CD45^+^PD-1^+^) and PD-L1^+^ tumor cells (EpCAM^+^PD-L1^+^) were elevated in the APOBEC tumors compared to control tumors, suggesting an active T cell-mediated immune response and potential feedback signaling leading to T cell dysfunction (Fig. 2E, F). A similar expansion of CD8^+^ T cell and CD103^+^ DC populations was observed in the tumor-draining lymph nodes from mice with APOBEC tumors (Supplementary Fig. S4A). The defect in APOBEC tumor growth and enhanced immune infiltration measured by flow cytometry was also observed in an independent experiment (Supplementary Fig. S4B, C). To extend these results to a second breast cancer cell line, we engineered inducible A3B expression in the mouse breast cancer line, EMT6, (Supplementary Fig. S4D-F). As with SMF tumors, EMT6 APOBEC tumors grew more slowly and had increased infiltration of leukocytes when implanted in syngeneic BALB/c mice (Supplementary Fig. S4G-I).

Next, to examine the relationship between immune infiltration and tumor size, we measured the correlation between immune cell abundance and tumor size at endpoint in SMF tumors. CD8^+^ T cells, CD4^+^ T cells, CD103^+^ DCs, and IFNγ^+^ cells were each negatively correlated with tumor size (Fig. 2G). This suggests that the adaptive immune response may mediate the growth defect observed in APOBEC tumors.

The localization of T cells in the TME is an important factor that influences tumor immunity and responses to immunotherapy (reviewed by (43)). T cells can be excluded from the tumor core and instead localize to the periphery in murine models and human tumors (44–47), and this exclusion may be one mechanism of immune suppression. Therefore, to assess T cell localization in APOBEC tumors, we performed immunohistochemical (IHC) staining for CD3 on an independent cohort of SMF tumors. CD3 staining was consistent with flow cytometry analyses and revealed an increase in the total number of T cells in APOBEC tumors. The T cells were most concentrated on the periphery of the APOBEC tumors, although importantly, significant levels of T cells also infiltrated the tumor core (Fig. 2H, I).

### The growth defect of APOBEC tumors is dependent on the A3B catalytic activity, not A3B protein expression

To discern whether the antitumor immune response in APOBEC tumors was due to the catalytic activity of A3B, and to rule out the possibility that expression of the human A3B protein in mouse cells may be immunogenic, we generated a catalytically inactive A3B mutant by site-directed mutagenesis of one of the A3B catalytic domains (E255Q). SMF cells were transduced with lentivirus expressing the A3B catalytic mutant to generate SMF-A3B^inactive^ cells. Dox treatment led to expression of catalytically-dead A3B in these cells, but there was no detectable increase in deaminase activity (Supplementary Fig. S5A-C). SMF-A3B^inactive^ cells were then injected into the mammary glands of immunocompetent wildtype mice on dox water to induce expression of the full-length, catalytically dead A3B protein. Tumors expressing catalytically dead A3B (SMF-A3B^inactive^ + dox) grew at similar rates and had similar numbers of total leukocytes (CD45^+^ cells) and T cells (CD3^+^ cells) as control tumors (Supplementary Fig. S5D, E). This indicates that the growth defect and immune response in APOBEC tumors is dependent on A3B catalytic activity and is not the result of expression of the human A3B protein in mouse cells.

To further explore whether the tumor growth defect and immune response of APOBEC tumors was due to A3B-mediated mutagenesis, as opposed to the expression of A3B protein, we took advantage of the reversibility of the dox-inducible system. SMF-A3B cells were cultured with dox in the cell medium for two weeks to mutagenize the cells and then dox was removed to downregulate A3B expression. These *in vitro* APOBEC mutagenized cells retain A3B-catalyzed mutations but do not express A3B protein. The proliferation rate of *in vitro* APOBEC mutagenized cells was similar to control, non-mutagenized cells (Supplementary Fig. S5F). In contrast, when implanted into the mammary gland of wildtype mice without dox in their drinking water (Supplementary Fig. S5G), the *in vitro* APOBEC mutagenized tumors grew more slowly than control tumors and had evidence of an increased adaptive immune response, as measured by qRT-PCR for T cell-specific genes *Gzma*, *Prf-1*, *Tbx21* (Supplementary Fig. S5H, I). The growth defect of *in vitro* APOBEC mutagenized tumors was not evident in NSG mice (Supplementary Fig. S5J), further confirming the role of the adaptive immune response in mediating the growth defect of APOBEC tumors. Together these data reveal that A3B activity promotes an infiltrated-inflamed TME in HER2-driven murine tumors and leads to an immune-dependent growth defect.

### APOBEC activity slows breast tumor growth by stimulating a tumor antigen-specific adaptive immune response

To understand the basis of the immune-mediated growth defect of APOBEC tumors, we performed RNA-sequencing on control and APOBEC tumors from either immunocompetent wildtype mice or immunodeficient NSG mice. APOBEC tumors in wildtype mice showed a significant upregulation of adaptive immune response gene ontology (GO) terms, including regulation of T cell mediated immunity/cytotoxicity/differentiation, antigen processing and presentation, and B cell activation (Fig. 3A, Supplementary Fig. S6A). Moreover, the top two pathways enriched in the APOBEC tumors in wildtype mice by gene set enrichment analysis (GSEA) were allograft rejection (Supplementary Fig. S6B) and IFNγ response (Fig. 3B), suggesting an adaptive immune response mechanism of tumor cell killing. In APOBEC tumors harvested from immunodeficient NSG mice, in contrast, the DNA repair pathway was significantly enriched by GSEA (Fig. 3B, Supplementary Fig. S6C), possibly due to the activation of repair pathways following the generation of A3B-catalyzed uracil lesions in the genome.

**Figure 3:**
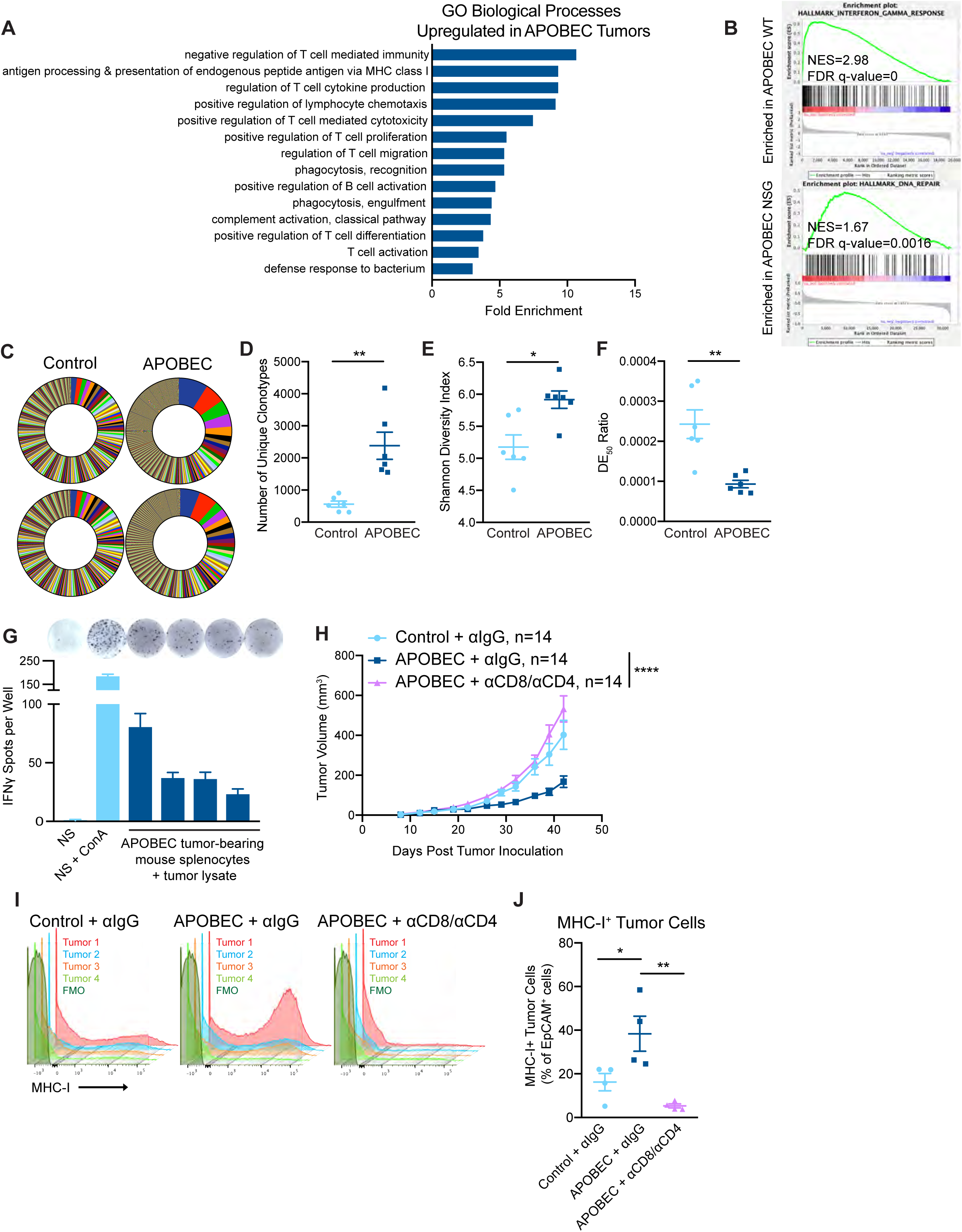
T cell-dependent antitumor responses in APOBEC tumors. **(A)** Gene ontology (GO) analysis of differentially expressed genes between control and APOBEC tumors in immunocompetent mice. Bar graph shows the fold enrichment of select GO biological processes that were significantly enriched in APOBEC tumors (n=6) compared to control tumors (n=6) (FDR<0.05, Fisher’s test). All significantly upregulated biological process GO terms are shown in Supplementary Fig. S6A. **(B)** Gene set enrichment analysis (GSEA) of differentially expressed genes between control and APOBEC tumors. Representative gene sets enriched in APOBEC tumors in immunocompetent mice (top) or immunodeficient mice (bottom) are shown. Normalized Enrichment Scores (NES) and FDR q-values are shown. All significantly enriched gene sets are shown in Supplementary Fig. S6B, C. **(C)** T cell receptor (TCR) sequencing from control (n=6) and APOBEC tumors (n=6) from wildtype mice. Pie charts show unique TCR clonotypes ranked by abundance in two control and two APOBEC tumors. **(D)** Quantification of the total number of unique clonotypes in control and APOBEC tumors (n=6 per cohort). **(E)** The Shannon diversity index of the TCR repertoire in control and APOBEC tumors. **(F)** TCR diversity evenness 50 (DE50) ratios in control and APOBEC tumors. DE50 ratio is calculated by the number of clonotypes composing the top 50% of total read counts divided by the total number of read counts. Error bars in (C-E) denote mean ± SEM. Statistical significance was determined by unpaired Student’s t test in (D) and unpaired Student’s t test with Welch’s correction in (C) and (E). **(G)** Representative ELISpot images and quantification of the number of IFNγ spots per well for each condition. NS, naïve splenocytes from a non-tumor-bearing mouse. ConA, concanavalin A model antigen. Error bars denote mean ± SD from 4 technical replicates per condition. **(H)** Tumor volume (mm3) over time for control tumors treated with isotype control antibody (n=14) and APOBEC tumors treated with isotype control (n=14) or αCD8 and αCD4 depletion antibodies (n=14) in wildtype mice. Error bars denote mean ± SEM. Statistical significance was determined by two-way repeated-measures ANOVA and Tukey’s multiple comparisons test. **(I)** Flow cytometry histograms showing MHC-I expression on EpCAM+ tumor cells from tumors in (H). **(J)** Quantification of MHC-I+ cells, expressed as a percentage of EpCAM+ cells, in tumors (n=4 per cohort) from (H). Error bars denote mean ± SEM. Statistical significance was determined by one-way ANOVA and Tukey’s multiple comparisons test. * p < 0.05, ** p < 0.01, **** p < 0.0001

Given that antigen presentation pathways were upregulated in APOBEC tumors, we were next interested in studying tumor-specific antigen responses in APOBEC tumors. To gain insight into these responses, we assessed changes in the T cell repertoire between control and APOBEC tumors using T cell receptor (TCR)-sequencing. RNA was extracted from control and APOBEC tumors growing in wildtype mice and used for TCR library preparation and sequencing of the β chain. The CDR3 variants were interrogated and unique clonotypes were counted. APOBEC tumors had more unique TCR clonotypes than control tumors (Fig. 3C, D). Using the Shannon entropy diversity index to measure the diversity richness of the clonotypes in the population, we found that APOBEC tumors had a higher clonotype diversity than control tumors (Fig. 3E). Finally, we used the diversity evenness 50 (DE_50_) ratio, which is a measure of the number of clonotypes making up the top 50% of reads relative to the total number of reads, to assess clonotype evenness. A high DE_50_ ratio indicates that clonotypes are evenly represented in the population, whereas a low DE_50_ ratio corresponds to a TCR repertoire that is dominated by specific CDR3 clonotypes. This analysis indicated that APOBEC tumors had a lower DE_50_ ratio than control tumors (Fig. 3F). Taken together, these analyses indicate that the TCR repertoire of APOBEC tumors exhibit increased diversity richness but decreased diversity evenness; interestingly, this pattern has been associated with productive T cell responses with antitumor effects and successful treatment with immunotherapy (48).

Tumor antigen-specific responses were examined by isolating splenocytes from APOBEC-tumor bearing mice and co-culturing the cells with autologous tumor cell lysate for 48 hours. Re-stimulation responses were measured by IFNγ ELISpot. Autologous APOBEC tumor lysate was capable of re-stimulating splenocytes from APOBEC-tumor bearing mice to produce IFNγ at comparable levels to that of naïve splenocytes (NS) stimulated with model antigen, concanavalin A (ConA; Fig. 3G). Thus, A3B-mediated mutagenesis may lead to the generation of tumor-specific antigens which are targeted by T cells.

### CD4^+^ T cells are required for the tumor growth defect of APOBEC tumors

To explore the requirement for T cells in mediating the antitumor immune response against APOBEC tumors, we depleted CD8^+^ T cells in APOBEC tumor-bearing mice using an anti-CD8 depleting antibody. We confirmed that CD8^+^ T cells were completely depleted in the peripheral blood using flow cytometry, and in the tumor at endpoint using qRT-PCR (Supplementary Fig. S7A-C). Interestingly, the growth defect of APOBEC tumors was not rescued upon CD8^+^ T cell depletion alone (Supplementary Fig. S7D). We next depleted CD8^+^ T cells and CD4^+^ T cells simultaneously (Supplementary Fig. S7E-G). In the absence of both CD4^+^ and CD8^+^ T cells, the APOBEC tumor growth defect was completely rescued, and APOBEC tumors grew similarly to the control tumors (Fig. 3H). Control or CD4/CD8-depleted tumors were harvested to assess major histocompatibility complex class I (MHC-I) expression on tumor cells. In the presence of T cells, APOBEC tumors had higher expression of MHC-I on tumor cells compared to control tumors. In contrast, when T cells were depleted, MHC-I expression on tumor cells was abrogated (Fig. 3I, J). Together, these results reveal that T cells are required for MHC-I upregulation and slowed tumor growth in APOBEC tumors.

### APOBEC activity renders murine HER2-driven breast tumors responsive to immune checkpoint inhibition

Because we found that A3B expression stimulated a T cell-mediated antitumor immune response, we next asked if APOBEC activity renders the tumors responsive to checkpoint inhibition. SMF-A3B cells were implanted in the mammary glands of wildtype mice and mice were administered dox water or control water. When control and APOBEC tumors reached 5 mm in diameter, mice were treated with combination anti-PD-1 and anti-CTLA-4 therapy twice weekly. Control tumors did not benefit from checkpoint inhibition, consistent with the clinical observation that checkpoint inhibition is not been effective in HER2^+^ breast cancer patients. In contrast, APOBEC tumor growth was significantly blunted upon treatment with anti-PD-1/anti-CTLA4 therapy (Fig. 4A). We defined a complete response (CR) as a full tumor regression (−100% change in tumor volume from the treatment start) and a partial response (PR) as any reduction in tumor volume from the treatment start. Checkpoint inhibitor treatment led to a partial response in only 1 of the 13 control tumors (Fig. 4B, Supplementary Fig. S8A). In contrast, 7 out of 11 APOBEC tumors had a complete or partial response to combination checkpoint inhibition (Fig. 4B, Supplementary Fig. S8A). Interestingly, both control and APOBEC tumors did not respond to anti-PD-1 monotherapy (Supplementary Fig. S8B). These results show that APOBEC activity sensitized HER2-driven murine breast cancers to combination anti-PD-1/anti-CTLA-4 checkpoint inhibition, but not single agent therapy.

**Figure 4:**
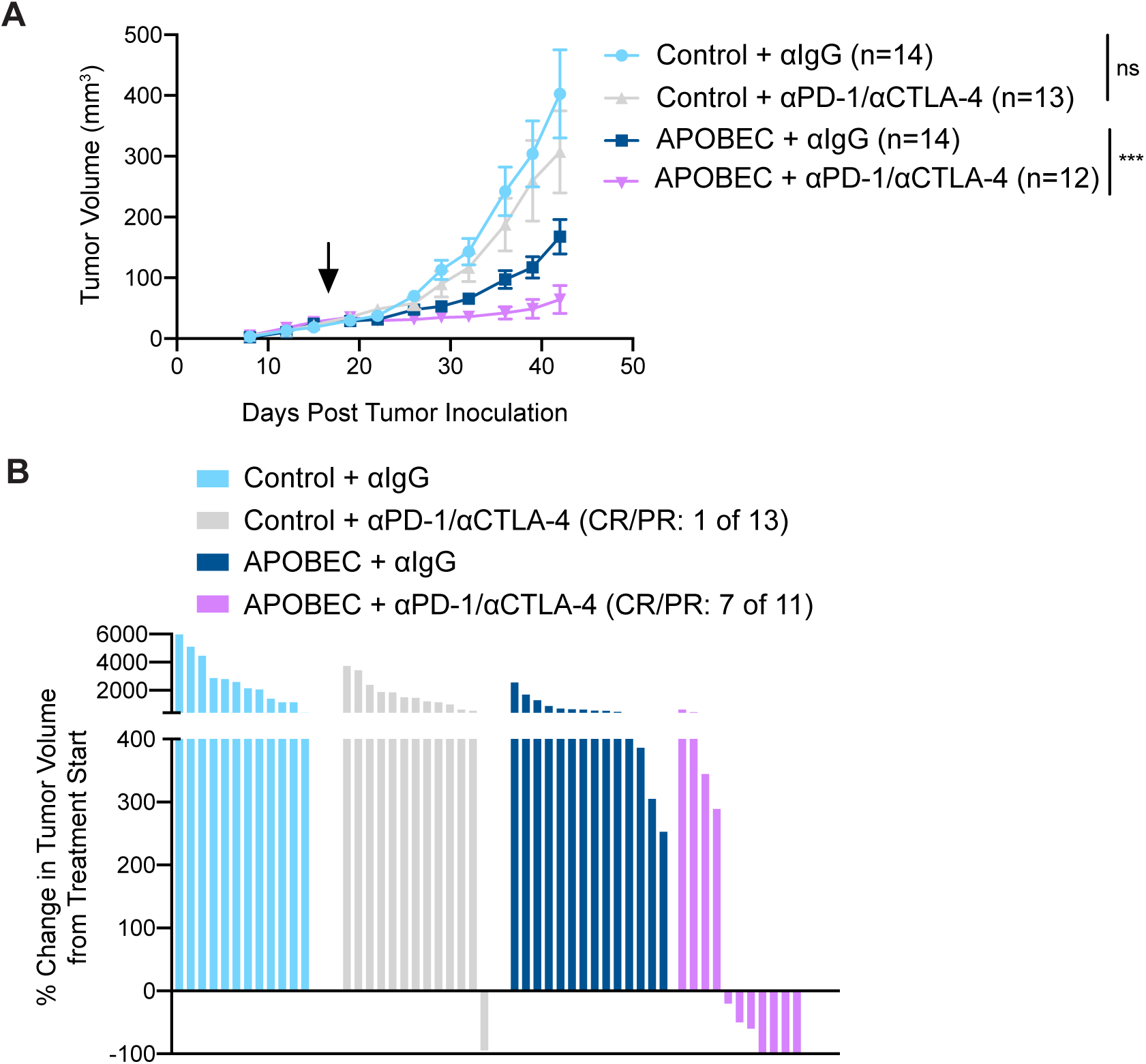
APOBEC tumors are sensitive to combination anti-PD-1/anti-CTLA4 immune checkpoint blockade. **(A)** Tumor volume (mm3) over time for control and APOBEC tumors treated with isotype control or αPD-1/αCTLA-4 antibodies. Arrow indicates the treatment start. Error bars denote mean ± SEM. Statistical significance was determined by two-way repeated-measures ANOVA and Tukey’s multiple comparisons test. **(B)** Change in tumor volume from treatment start for palpable tumors until endpoint. Each bar denotes an individual tumor. CR, complete response; PR, partial response. ns p > 0.05, *** p < 0.001

### APOBEC-mediated genetic heterogeneity permits immune escape, while clonal APOBEC tumors remain in cancer-immune equilibrium

Genomic studies of human cancer suggest that episodic APOBEC mutagenesis may fuel cancer heterogeneity and evolution (49–51). In melanoma and NSCLC, mutational and neoantigen heterogeneity reduces antitumor immunity (52–54) and response to checkpoint inhibitor therapy. For instance, lung tumors with more clonal neoantigens are better controlled by neoantigen-specific T cells and have improved responses to checkpoint inhibitors (55). The role of intratumor diversity in breast cancer immunogenicity has yet to be thoroughly studied. Thus, we were interested in understanding the consequences of APOBEC-mediated genetic heterogeneity on antitumor immunity and mammary tumor growth in immunocompetent mice.

To assess differences between heterogenous and clonal APOBEC tumors, SMF-A3B cells were cultured with dox for 2 weeks to mutagenize and induce genetic heterogeneity in the population of cells, and then dox was removed to downregulate A3B. These cells are referred to as “parental APOBEC”, whereas the control, non-mutagenized cells are referred to as “parental control”. We next derived single-cell clones by limiting dilution from the parental APOBEC and parental control populations. We screened several clonal populations for the ability to grow at the same rate as parental populations *in vitro*. Control clone 1 and APOBEC clone 1 grew slower than the parentals, while control clone 2 and APOBEC clone 2 grew at the same rate as parentals (Fig. 5A). When we implanted the clones in immunocompromised, athymic nude mice and measured tumor growth, only control clone 2 and APOBEC clone 2 were able to form tumors similarly to the parental counterparts (Fig. 5B). Therefore, we proceeded to study the tumor growth of control clone 2 and APOBEC clone 2 in immunocompetent, wildtype mice. While control clone 2 grew similarly to the parentals, APOBEC clone 2 cells gave rise to very small tumors that remained in a cancer-immune equilibrium until the animals were sacrificed (Fig. 5C). When we compared the average size of tumors formed in the presence or absence of the adaptive immune response, we found tumors formed from APOBEC clone 2 were significantly smaller in wildtype mice (Fig. 5D). Thus, APOBEC-mediated heterogeneity may limit the potential of a fully productive immune response against hypermutated breast tumors. In contrast, clonal APOBEC tumor growth may be controlled in cancer-immune equilibrium.

**Figure 5:**
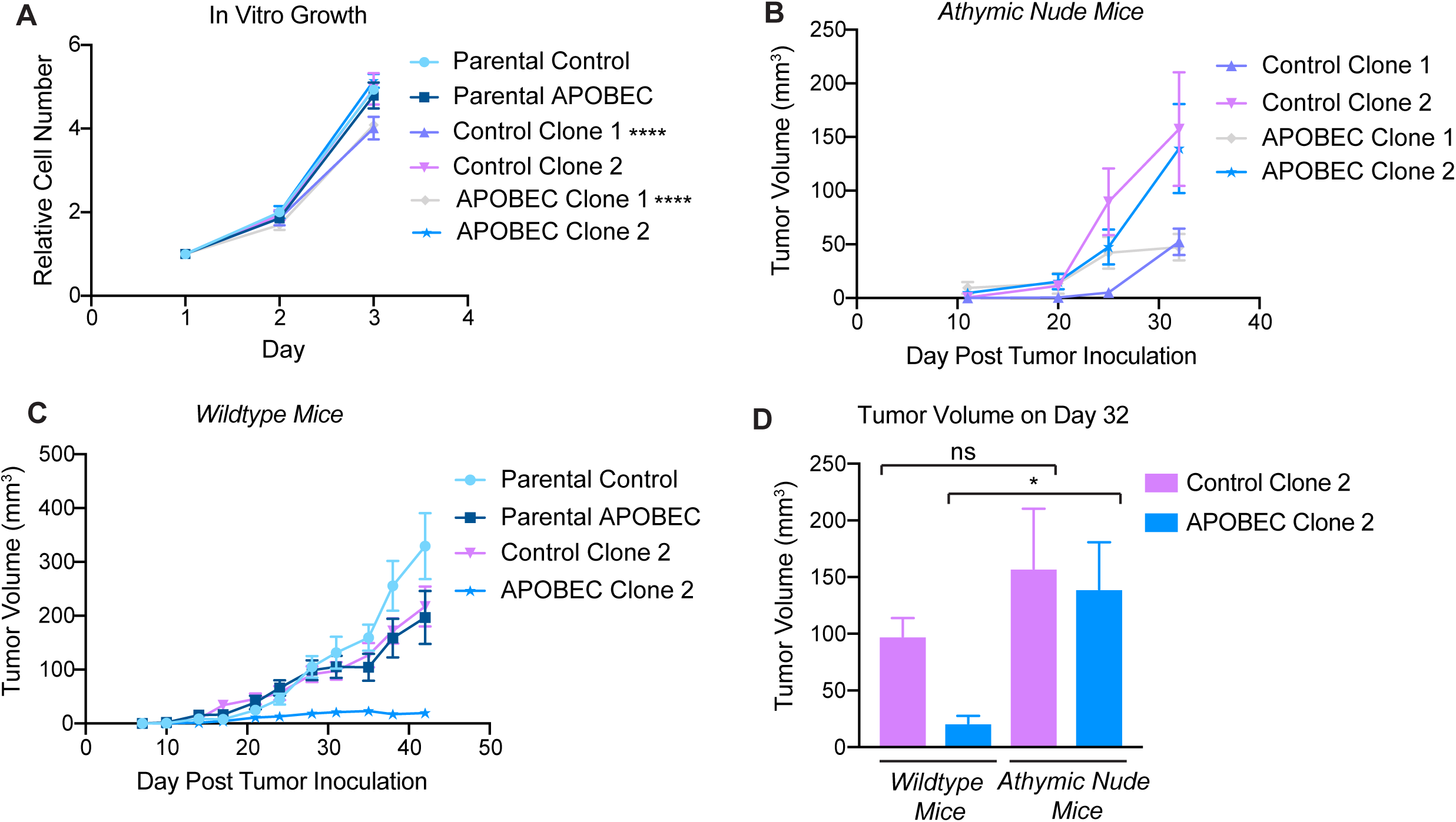
Complete immune-mediated suppression of clonal APOBEC tumor growth. **(A)** In vitro growth curves of single-cell clones derived from control SMF-A3B cells (control) or SMF-A3B cells treated with dox for 2 weeks (APOBEC). The growth curves of the polyclonal parental cells (parental control or parental APOBEC) are shown as a control. Error bars denote mean ± SD of 4 replicates. Control Clone 1 and APOBEC Clone 1 proliferate more slowly than the parental control cells, as determined by two-way ANOVA and Dunnett’s multiple comparison test. **(B)** Tumor volume (mm3) over time for control and APOBEC clones injected in the mammary gland of athymic nude mice. Error bars denote mean ± SEM. **(C)** Tumor volume (mm3) over time for control and APOBEC clones, as well as the corresponding polyclonal parental populations, injected in the mammary gland of immunocompetent wildtype mice. Error bars denote mean ± SEM. **(D)** Comparison of tumor volume on day 32 between clones grown in immunocompromised nude mice from (B) and immunocompetent wildtype mice from (C). Error bars denote mean ± SEM and statistical significance was determined by one-way ANOVA and Sidak’s multiple comparisons test. ns p > 0.05, * p < 0.05, **** p < 0.0001

### The APOBEC mutational signature is associated with an adaptive immune response in basal-like but not HER2-enriched human breast cancers

We were next interested in determining whether human breast tumors with APOBEC mutagenesis have evidence of an increased adaptive immune response. To do this, we analyzed breast tumors from TCGA for which both whole-exome sequencing (WES) and RNA-sequencing data were available. To assess the enrichment of APOBEC mutational signatures, we analyzed WES data using an established algorithm that quantifies the enrichment of C-to-T or C-to-G mutations occurring in the T<u>C</u>W context relative to all other cytosine mutations (9) (Fig. 6A). Similar to previous reports (2,9,12), we found that the HER2-enriched subtype had the highest median APOBEC enrichments scores and the largest proportion of tumors with enrichment scores > 2 (Fig. 6A). To estimate immune cell infiltration, we analyzed RNA-seq data for the expression of individual immune checkpoint genes or immune cell gene signatures (56, 57) (Supplementary Table 1). In this manner, we were able to generate quantitative estimates of APOBEC mutagenesis and immune cell infiltration within individual tumors (Supplementary Table 2). We first examined the relationship between APOBEC mutagenesis and the expression of immune signatures in HER2-enriched and basal-like breast cancers as determined by the PAM50 subtype. The basal-like category includes most of the TNBCs and is considered the most immunologically active subtype of breast cancer (58, 59). We segregated basal-like and HER2-enriched tumors into APOBEC-high (Fig. 6B) or APOBEC-low groups (Supplementary Fig. S9A) using an APOBEC enrichment score cutoff of 2 (60). Hierarchical clustering of tumors based on immune cell signatures revealed two main clusters in each subtype. Tumors in cluster 1 had high expression of immune signatures that were reflective of an antitumor adaptive immune response, including type-1 T helper cells (Th1 cells), activated DCs (aDCs), CD8^+^ T cells, cytotoxic cells, interferon signaling pathway (IFN), major histocompatibility complex class II antigen presentation pathway (MHC-II), and checkpoint genes such as *LAG3, PD1, PDL1, PDL2, CTLA4,* and *TIM3*. Tumors in cluster 2 had low expression of antitumor immune response signatures and high expression of several immunosuppressive gene signatures, such as macrophages and neutrophils.

**Figure 6:**
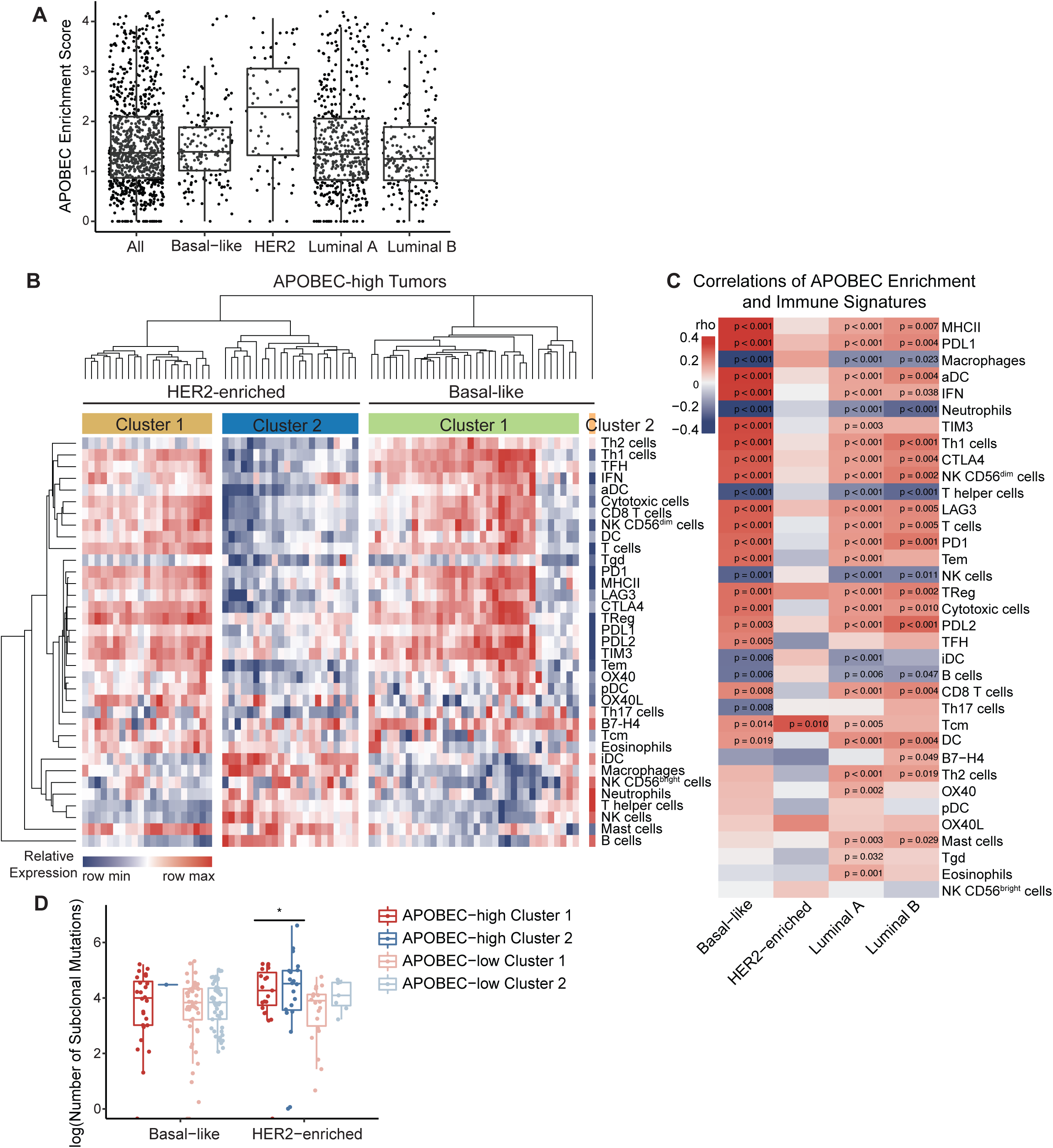
The TME phenotype of APOBEC-high human breast cancers is dependent on molecular subtype and the number of subclonal mutations. **(A)** APOBEC enrichment score calculated from whole-exome sequencing (WES) data for all TCGA breast cancer samples. APOBEC enrichment score of 2 or higher delineates APOBEC-high tumors. Boxplots show 25th percentile, median, and 75th percentile, while whiskers show minimum and maximum values excluding outliers. **(B)** Heatmap showing the relative expression of immune cell gene signatures from TCGA RNA-seq data in APOBEC-high tumors, grouped by breast cancer subtype. Columns are individual patient tumors and rows are different immune cell gene signatures. Legend shows colors corresponding to relative expression levels (red, row max; blue, row min). Hierarchical clustering segregated tumors into 2 main clusters in the HER2-enriched subtype and 2 clusters in the basal-like subtype. **(C)** Heatmap showing correlation Spearman rho values between APOBEC enrichment score and immune gene signatures for each molecular subtype of breast cancer. p-values are shown and legend shows colors corresponding to rho value (red, immune signature positively correlated with APOBEC enrichment score; blue, immune signature negatively correlated with APOBEC enrichment score). **(D)** The number of subclonal mutations (from Raynaud et al.) in APOBEC-high clusters depicted in (B) and APOBEC-low clusters depicted in Supplementary Figure S9. APOBEC-high HER2-enriched tumors in cluster 2 had more subclonal mutations than tumors in cluster 1. Boxplots show 25th percentile, median, and 75th percentile, while whiskers show minimum to maximum values excluding outliers. Statistical significance was determined one-way ANOVA and Sidak’s multiple comparisons test. * p < 0.05. MHC-II, major histocompatibility complex class II antigen presentation; IFN, interferon signaling pathway; Th1 cells, type-1 T helper cells; Th2 cells, type-2 T helper cells; Tgd, T gamma delta cells; Treg, T regulatory cells; Tem, T effector memory cells; Tcm, T central memory cells; TFH, T follicular helper cells; Th17, T helper 17 cells; DC, dendritic cells; aDC, activated dendritic cells; iDC, immature dendritic cells; pDC, plasmacytoid dendritic cells; CD56dim NK cells, CD56 dim natural killer cells; CD56bright NK cells, CD56 bright natural killer cells.

Nearly all of the APOBEC-high basal-like tumors fell within cluster 1, reflective of an antitumor adaptive immune response (Fig. 6B). These results are consistent with the well-defined hot TME of TNBC and their response to immune checkpoint inhibition. Surprisingly, in contrast to basal-like tumors, half of the APOBEC-high HER2-enriched tumors fell within cluster 1 and half within cluster 2 (Fig. 6B).

To further explore the differences in immune cell gene expression signatures between basal-like, HER2-enriched, and luminal A/B tumors, we analyzed the correlation between the APOBEC signature enrichment score and each immune cell gene signature, as measured by a quantitative score (Supplementary Table 3). This analysis showed that the correlation between APOBEC signatures and immune cell infiltration varied by breast cancer subtype. In the basal-like subtype, APOBEC signature enrichment score was positively correlated with numerous adaptive immune response gene signatures (e.g. MHC-II, aDCs, IFN, Th1 cells) and checkpoint genes (e.g. *PDL1*, *TIM3*, *CTLA4*), and negatively correlated with known immunosuppressive cell types (macrophages, neutrophils) (Fig. 6C). Luminal A and B subtypes showed similar patterns of correlation between APOBEC enrichment and immune signatures (Fig. 6C). In contrast, all but one of the immune signatures (Tcm, T central memory cells), did not significantly correlate with APOBEC enrichment score in the HER2-enriched subtype, despite this subtype possessing the highest median APOBEC enrichment scores (Fig. 6C). In summary, the APOBEC mutational signature is associated with antitumor adaptive immunity gene expression in basal-like breast cancer patients, but there was no evidence of association in HER2-enriched patients.

### APOBEC-high HER2-enriched tumors in cluster 2 have increased subclonal mutations compared to tumors in cluster 1

To understand the differences in immune infiltration between APOBEC-high HER2-enriched tumors in cluster 1 and 2, we first examined clinical features of tumors from each cluster. There were no statistically significant differences in estrogen receptor status, p53 status by IHC, node positivity, risk of recurrence, or pathological stage between cluster 1 and 2 of APOBEC-high HER2 tumors (data not shown). Next, in light of previous findings that tumors with more subclonal mutations have a less productive immune response (52–55), we postulated that genetic heterogeneity may underly the TME differences between APOBEC-high basal-like and HER2-enriched tumors. To test this, we used the clonal phylogenies of TCGA breast cancers generated by Raynaud and colleagues (61) to explore the relationship between subclonal mutations and immunogenicity in human breast cancer. The HER2-enriched subtype is characterized as the breast cancer subtype with the highest levels of intratumor heterogeneity, as measured by number of clones in the tumor phylogeny (61). Further, HER2^+^ breast cancers have increased allelic imbalance and chromosomal instability compared to HER2-negative tumors (62).

We compared the number of subclonal mutations between tumors with a hot TME (cluster 1) and tumors with a cold TME (cluster 2). Interestingly, in HER2-enriched tumors, APOBEC-high tumors in cluster 2 had more subclonal mutations than APOBEC-high tumors in cluster 1 (Fig. 6D), despite the fact that the APOBEC enrichment scores were similar between these two groups (Supplementary Fig. S9B). This suggests that the immunogenicity of APOBEC-high tumors between breast cancer subtypes may be due to the levels of intratumor genetic diversity. Basal-like tumors are less heterogenous and have lower APOBEC enrichment scores on average than HER2-enriched tumors. Conversely, HER2-enriched tumors with high APOBEC enrichment scores and high levels of genetical heterogeneity may undergo immune escape and acquire a cold TME.

## DISCUSSION

APOBEC mutational signatures have been identified in more than 22 different cancer types (9, 10), but the functional consequences of APOBEC activity on the tumor immune microenvironment have not been explored. Here we show that APOBEC activity promotes an immunologically hot, infiltrated-inflamed tumor microenvironment, leading to slowed tumor growth. We find that the slowed growth of APOBEC tumors is due to an adaptive immune-mediated mechanism that requires the activity of CD4^+^ T cells. APOBEC tumors exhibit a T cell-dependent upregulation of MHC-I expression on tumor cells, and this is associated with increased TCR diversity within tumors. Consistent with increased immune cell infiltration, APOBEC tumors are sensitive to checkpoint inhibitors. While other studies have examined how APOBEC mutagenesis sensitizes tumors to checkpoint inhibitors, this is the first study to our knowledge to comprehensively define the direct consequences of APOBEC activity on the tumor immune microenvironment in the absence of therapy.

The role of CD4^+^ T cells in the APOBEC-dependent antitumor immune response is intriguing and opens up the possibility for CD4^+^ T cell-directed therapies, such as CTLA-4 inhibitors or CD4^+^ T cell adoptive transfer, to treat APOBEC-high patients. In a recent study of murine APOBEC3-mutagenized models of TNBC, the function of T follicular helper cells in activating B cells and antibody generation was found to be required for sensitivity to anti-CTLA-4/anti-PD-1 checkpoint blockade (31). Our work shows that A3B activity sensitizes HER2-driven mammary tumors to anti-CTLA-4/anti-PD-1 combination therapy, while anti-PD-1 monotherapy alone was ineffective. Similarly, Hollern and colleagues found that single-agent anti-PD-1 was inferior to the combination therapy for TNBC (31). Thus, while the majority of immunotherapy trials focus on re-invigorating CD8^+^ cytotoxic T cells, our findings and others suggest that harnessing the activity of CD4^+^ helper T cells may be more beneficial for breast tumors with APOBEC mutational signatures.

At the same time that APOBEC-catalyzed mutations may promote immunogenicity, APOBEC activity can also generate genetic heterogeneity and fuel tumor evolution (7, 63). For example, extensive evidence of APOBEC mutagenesis was found in lung cancers harboring the highest burden of subclonal mutations (50), and more than 45% of subclonal mutations in cancer genes could be explained by APOBEC mutagenesis (51). While there is growing interest in understanding how intratumor genetic diversity impacts productive immune responses, little is known about the effects of APOBEC-catalyzed subclonal diversification on tumor immunogenicity. When we examined the relationship between APOBEC mutagenesis and immunogenicity in human breast cancers, we observed a strong correlation between APOBEC enrichment scores and immune cell gene signatures in basal-like tumors, consistent with findings in other tumor types (60,64,65). In contrast, there was no correlation between APOBEC enrichment and immune cell signatures in HER2-enriched breast cancers. In fact, half of HER2-enriched tumors with high APOBEC enrichment scores (cluster 2) had low expression of adaptive immune signatures. At first glance, this was a surprising result – especially in light of our finding that APOBEC activity promotes immune infiltration in HER2-driven mouse mammary tumors – and suggested that cluster 2 tumors may have evolved immune-suppression mechanisms that limit an antitumor adaptive immune response. While the details of such mechanisms remain unknown, initial insight came from examining the frequency of subclonal mutations in these tumors. Among APOBEC-high tumors, immune-suppressed (cluster 2) tumors had a higher number of subclonal mutations than immune-infiltrated (cluster 1) tumors. These results are reminiscent of findings from other groups. For instance, in lung cancer, high clonal neoantigen burden is associated with neoantigen-reactive T cells and improved immunotherapy response (55). In breast cancer, tumors with high levels of heterogeneity have less infiltration of antitumor immune cells, including CD8^+^ and CD4^+^ T cells, lower expression of PD-L1, and lower expression of cytolytic enzymes, granzyme A and perforin-1 (66). These results suggest that a subset of APOBEC-high HER2 tumors with a high frequency of subclonal mutations can evade immune activation. These results mirror our findings in mouse tumors, where clonal APOBEC tumors are controlled by the immune system more profoundly than polyclonal APOBEC tumors. We propose a model (Supplementary Fig. S9C), where APOBEC mutagenesis leads to immune infiltration and immunotherapy benefit in both mouse models and human breast tumors yet can also foster subclonal diversification to promote evasion of the immune response.

Therefore, to exploit the immunogenic nature of APOBEC mutations without allowing acceleration of the aggressiveness of the tumor, immunotherapy could be used early on to target the subclones already harboring APOBEC-catalyzed neoantigens and prevent further diversification. In fact, clinical trials of immunotherapy in breast cancer show that tumors respond better when administered in earlier lines of therapy (reviewed by (20)). Given our findings, prior evidence in murine models (31, 32), and human genomic studies implicating APOBEC mutagenesis in immune infiltration (60,64,65) and immunotherapy response (27–30,67), endogenous APOBEC mutagenesis may render human tumors responsive to immunotherapy. Thus, APOBEC mutational signatures and mutational clonality may be useful biomarkers predicting response to immunotherapy in women with breast cancer. This is particularly notable for HER2^+^ breast cancer, because while the majority of reports of durable clinical benefit and newly initiated immunotherapy trials are for TNBC (reviewed by (20)), HER2^+^ breast cancers have the highest median levels of APOBEC enrichment compared to other breast cancer subtypes.

Finally, our findings that APOBEC activity slows the growth of polyclonal tumors through an antitumor immune-mediated response are in contrast to a recent study of another mutational process, showing that UVB-derived mutational heterogeneity reduces antitumor immunity and generates highly aggressive tumors that grow faster than non-mutagenized tumors (54). Interestingly, the UVB mutational signature does not predict response to checkpoint blockade in melanoma patients (29). This raises the possibility that APOBEC-mediated mutagenesis is a particularly immunogenic mutational process, compared to other mutagens, such as UVB irradiation, or a general increase in the TMB. For instance, APOBEC SBS signature 13, but not overall TMB, correlates with immune response-specific gene expression in breast cancer (64). Additionally, the APOBEC mutational signature is a better predictor of durable clinical benefit to immunotherapy than total TMB in NSCLC (28). Lastly, in a cohort of patients with diverse cancer types, APOBEC signatures correlate with improved immunotherapy response, independent of TMB (27). It is possible that APOBEC-mediated mutations generate neoantigens that are particularly immunogenic (e.g. with increased hydrophobicity (27)) or are more likely to occur in highly expressed genes or regions of open chromatin (e.g. R-loops (68, 69)), although human data shows an inverse correlation between C-to-T mutations and gene expression (1). Future work should focus on the mechanism by which the APOBEC mutational process generates immunogenic neoantigens.

## MATERIALS AND METHODS

### Tissue culture and reagents

All cell lines were grown at 37°C in 5% CO_2_. SMF cells were provided by Dr. Lewis Chodosh (University of Pennsylvania) and were cultured in Dulbecco’s Modified Eagle Medium (DMEM), 10% fetal bovine serum (FBS), 1% L-glutamine (Gibco 25030-081), 1% penicillin/streptomycin (Gibco 15140-122), and 5 µg/mL insulin (Gemini Bioproducts 700-112P). EMT6 cells were provided by the Duke Cell Culture Facility and were cultured in Waymouth’s Medium 752/1, 15% FBS, 1% L-glutamine, and 1% penicillin/streptomycin. BT474 cells were cultured in RPMI-1640, 10% FBS, 1% L-glutamine, and 1% penicillin/streptomycin. SKBR3 cells were cultured in DMEM, 10% FBS, 1% L-glutamine, and 1% penicillin/streptomycin. SMF-A3B cells were selected in 1 µg/mL puromycin (Sigma P8833-10MG) and 1 mg/mL neomycin (G418, Sigma, 345810-1GM). EMT6-A3B cells were selected in 4 µg/mL puromycin and 1 mg/mL neomycin. Doxycycline (RPI D43020-100.0) was added to the cell medium to induce the expression of A3B where specified at concentrations described. Cells were harvested for qRT-PCR, deaminase activity assay, or Western blot analysis.

Cell viability assays were performed using CellTiter-Glo (Promega) according to manufacturer instructions. Cells were plated at 2,000 cells per well in an opaque 96-well plate and treated with doxycycline on day 0. Doxycycline in cell medium was refreshed every 3 days.

Colony formation assays were performed by plating cells at 2,000 cells per 10-cm plate and cultured with doxycycline for 14 days. Doxycycline in cell medium was refreshed every 3 days. PBS was used to wash plates and 0.5% crystal violet in 25% methanal was used to stain cell colonies for 5 mins. The plates were dried overnight and imaged. Colonies were quantified using ImageJ Fiji.

For immunofluorescence staining of adherent cells, 5×10^4^ cells per well were plated on coverslips with 0.1% gelatin in a 24-well plate. 1 µg/mL doxycycline was added, and cells were cultured for 3 days prior to fixation in 4% paraformaldehyde. Coverslips were washed in PBS, permeabilized in 0.5% Triton-X 100, washed in PBS, and blocked in 3% BSA and 10% normal goat serum for 1 hour at room temperature. Coverslips were incubated with 1:800 HA-tag rabbit (Cell Signaling 3724S) primary antibody overnight at 4°C, washed, and incubated in with 1:500 goat anti-rabbit AF488 (Life Technologies A1103) secondary antibody for 1 hour at room temperature. Coverslips were then washed in PBS, stained with DAPI for 10 minutes, and mounted on slides with Prolong Gold (Thermo P36930). Slides were imaged Zeiss Axio Imager Widefield fluorescence microscope.

### Plasmids and viral transduction

To generate dox-inducible A3B expression in murine cancer cell lines, a 2-vector system was utilized. pLVX-Tet-On Advanced plasmid containing the rtTA cassette was provided by Dr. Ann Marie Pendergast (Duke University). pLenti-Tet-On-A3B plasmid containing tetracycline responsive human APOBEC3B gene (NM_004900.4) that is HA-tagged on the C-terminus was generated by VectorBuilder. The APOBEC3B gene contains an in-frame 66 bp SV40 T-antigen intron sequence to disrupt transcription of the gene in E. coli for successful cloning without introducing A3B-catalyzed mutations in the construct sequence. To generate the catalytically inactive mutant of A3B (E255Q), site-directed mutagenesis of the pLenti-Tet-On-A3B plasmid was performed by Genewiz. HEK293T cells were transfected with psPAX2 and pMDG.2 packaging plasmids (gifts from Didier Trono, EPFL, Lausanne, Switzerland; Addgene plasmids 12559 and 12660), the lentiviral expression plasmid, PLUS reagent (Thermo 11514015), and Lipofectamine 2000 (Thermo 11668019). 0.8 mM sodium butyrate was added to cell medium 1- and 2-days post-transfection to prevent epigenetic silencing of the lentiviral vector. Lentivirus was collected in the supernatant and filtered prior to concentrating with Lenti-X™ Concentrator (Clontech 631231) manufacturer protocol.

To generate SMF-A3B and EMT6-A3B cell lines, SMF cells and EMT6 cells were transduced at 50% confluency in 6-well plates with 1 mL of concentrated lentivirus and 6 µg/mL polybrene (Sigma 107689) at 1000xg and 33°C for 2 hours. Cells transduced with pLVX-Tet-On Advanced lentivirus were selected in neomycin for at least 10 days. Cells were then transduced with pLenti-Tet-On-A3B lentivirus and selected in puromycin for an additional 14 days.

### Animal work

Animal care and animal experiments were performed with the approval of, and in accordance with, guidelines of the Duke University IACUC. Mice were housed under barrier conditions with 12-hour light/12-hour dark cycles. Female FVB mice (FVB/NJ; used with SMF cells) and female BALB/c mice (BALBc/J; used with EMT6 cells) were obtained from The Jackson Laboratory. Female outbred athymic nude mice (J:NU) and female NOD.Cg-*Prkdc^scid^ Il2rg^tm1Wjl^*/SzJ (NSG) mice were obtained from The Jackson Laboratory.

Tumor cell lines were implanted in bilateral 4^th^ inguinal mammary fat pads of 6-8 week old female recipient mice. 2×10^6^ SMF-A3B cells or 2×10^4^ EMT6-A3B cells in complete cell medium were used for implantation. Tumors were monitored for growth, measured using calipers 2-3 times per week, and sacrificed at experimental endpoint or when tumors reached 10-15 mm in diameter. Tumor volume was calculated using (L*W*W*π)/6, where L is length of the longer side and W is length of the shorter side. Where indicated, 1 mg/mL of doxycycline supplemented with 5% sucrose was added to mouse drinking water 2 days prior to tumor cell implantation.

*In vivo* depletion antibodies were administered via intraperitoneal injected on day −2 and - 1 prior to implantation, then continued twice weekly until endpoint. 300 µg of anti-CD8 (BioXCell BE0117), or 300 µg of anti-IgG2b isotype control (BioXCell BE0090), was used for CD8 depletion alone. 200 µg of anti-CD8 (BioXCell BE0117) and 200 µg of anti-CD4 (BioXCell BE0003-1), or 400 µg of anti-IgG2b isotype control (BioXCell BE0090), was used for CD8/CD4 dual depletion. For anti-PD-1 monotherapy, antibodies were administered when the majority of tumors reached 5 mm in diameter, for a total of 3 doses in one week (day 13, 15, 17) and 3 doses in the next week (day 20, 22, 24). 200 µg of anti-PD-1 (BioXCell BE0146), or 300 µg of anti-IgG2b (BioXCell BE0090), was used for monotherapy. For combination anti-PD-1/anti-CTLA-4 therapy, antibodies were administered when the majority of tumors reached 5 mm in diameter and continued twice weekly until endpoint. 200 µg of anti-PD-1 (BioXCell BE0146) and 200 µg of anti-CTLA-4 (BioXCell BE0164), or 400 µg of anti-IgG2b isotype control (BioXCell BE0090), was used.

### Flow cytometry

Bilateral tumors were harvested and aggreged for each mouse, then minced into small chunks. Tumor chunks were digested with warmed digestion buffer containing 300 U/mL collagenase (StemCell 554656) and 100 U/mL hyaluronidase (StemCell 554656) at 37 °C for 1 hour, vortexing every 15 minutes. Digested tumors were incubated in ACK lysis buffer for 5 minutes to lysis red blood cells. Tumors were centrifuged, washed in stain buffer (BD Biosciences 554656), decanted, and resuspended in Dispase II (5 mg/mL; StemCell 7913) and DNase I (100 μg/mL; Worthington Biochemical LS002006) for 5 minutes, mixing. Tumors were then passed through 70 µm strainer, washed in stain buffer, counted, and 1×10^6^ cells in 100 µL of stain buffer were added to 96-well untreated v-bottom plate for staining. Prior to intracellular antigen staining, cells were activated using 2 µL of leukocyte activation cocktail with GolgiPlug (BD Biosciences 550583) for 3 hours at 37°C and 5% CO_2_.

LIVE/DEAD™ Fixable Aqua Dead Cell Stain Kit (Thermo L34957) was used to stain dead cells in PBS according to manufacturer protocol for 30 minutes at 4°C in the dark. Cells were washed in PBS three times and resuspended in 100 µL of PBS for antibody surface staining. 2 µL of CD16/CD32 Fc Block antibody (BD Biosciences 553141) was added for 10 minutes at 4°C in the dark. Surface antigen antibodies were added at dilutions listed below and incubated for 30 minutes at 4°C in the dark. Cells were washed in PBS and transferred to falcon tubes for analysis.

For intracellular antigen staining, cells were fixed in either Foxp3 fixation buffer (BD Biosciences 560409) or BD Cytofix^TM^ Fixation Buffer (BD Biosciences 554655) for 30 minutes at 4°C in the dark. Cells were washed and stored at 4°C in the dark overnight. Cells were permeabilizated in either Foxp3 permeabilization buffer (BD Biosciences 560409) for 30 minutes at 37°C or BD perm/wash buffer for 15 minutes at 4°C. Cells were washed in PBS and resuspended in 100 µL of PBS for intracellular antigen staining using the antibody dilutions listed below and incubated for 25 minutes at room temperature in the dark. Cells were washed in PBS and transferred to falcon tubes for analysis.

Cells were analyzed using a FACSCanto analyzer (BD Biosciences) and data were analyzed using FlowJo software (TreeStar, Ashland, OR). Fluorescence minus one (FMO; all antibodies in the panel, except for one) was used to determine proper gating of individual cell types. Individual cell type compartments were represented as either the percentage of total CD45^+^ cells or the percentage of total live cells. Treg and Th2 cell compartments were represented as percentage of total CD4^+^ T cells. The correlation between immune cell frequency and tumor volume was calculated using the mean volume of bilateral tumors at endpoint.

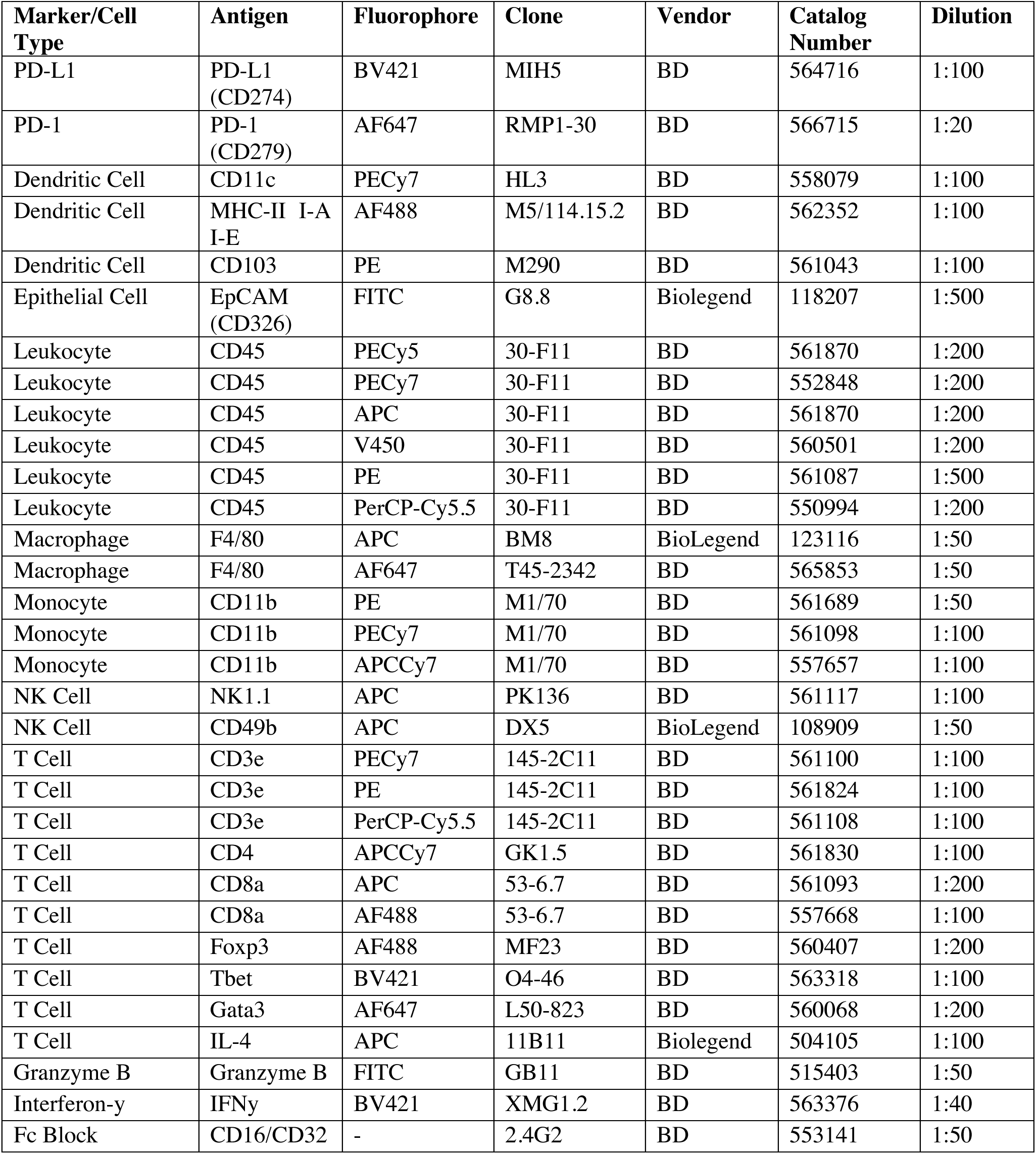

| Marker/Cell Type | Antigen | Fluorophore | Clone | Vendor | Catalog Number | Dilution |
| --- | --- | --- | --- | --- | --- | --- |
| PD-L1 | PD-L1 (CD274) | BV421 | MIH5 | BD | 564716 | 1:100 |
| PD-1 | PD-1 (CD279) | AF647 | RMP1-30 | BD | 566715 | 1:20 |
| Dendritic Cell | CD11c | PECy7 | HL3 | BD | 558079 | 1:100 |
| Dendritic Cell | MHC-II I-A I-E | AF488 | M5/114.15.2 | BD | 562352 | 1:100 |
| Dendritic Cell | CD103 | PE | M290 | BD | 561043 | 1:100 |
| Epithelial Cell | EpCAM (CD326) | FITC | G8.8 | Biolegend | 118207 | 1:500 |
| Leukocyte | CD45 | PECy5 | 30-F11 | BD | 561870 | 1:200 |
| Leukocyte | CD45 | PECy7 | 30-F11 | BD | 552848 | 1:200 |
| Leukocyte | CD45 | APC | 30-F11 | BD | 561870 | 1:200 |
| Leukocyte | CD45 | V450 | 30-F11 | BD | 560501 | 1:200 |
| Leukocyte | CD45 | PE | 30-F11 | BD | 561087 | 1:500 |
| Leukocyte | CD45 | PerCP-Cy5.5 | 30-F11 | BD | 550994 | 1:200 |
| Macrophage | F4/80 | APC | BM8 | BioLegend | 123116 | 1:50 |
| Macrophage | F4/80 | AF647 | T45-2342 | BD | 565853 | 1:50 |
| Monocyte | CD11b | PE | M1/70 | BD | 561689 | 1:50 |
| Monocyte | CD11b | PECy7 | M1/70 | BD | 561098 | 1:100 |
| Monocyte | CD11b | APCCy7 | M1/70 | BD | 557657 | 1:100 |
| NK Cell | NK1.1 | APC | PK136 | BD | 561117 | 1:100 |
| NK Cell | CD49b | APC | DX5 | BioLegend | 108909 | 1:50 |
| T Cell | CD3e | PECy7 | 145-2C11 | BD | 561100 | 1:100 |
| T Cell | CD3e | PE | 145-2C11 | BD | 561824 | 1:100 |
| T Cell | CD3e | PerCP-Cy5.5 | 145-2C11 | BD | 561108 | 1:100 |
| T Cell | CD4 | APCCy7 | GK1.5 | BD | 561830 | 1:100 |
| T Cell | CD8a | APC | 53-6.7 | BD | 561093 | 1:200 |
| T Cell | CD8a | AF488 | 53-6.7 | BD | 557668 | 1:100 |
| T Cell | Foxp3 | AF488 | MF23 | BD | 560407 | 1:200 |
| T Cell | Tbet | BV421 | O4-46 | BD | 563318 | 1:100 |
| T Cell | Gata3 | AF647 | L50-823 | BD | 560068 | 1:200 |
| T Cell | IL-4 | APC | 11B11 | Biolegend | 504105 | 1:100 |
| Granzyme B | Granzyme B | FITC | GB11 | BD | 515403 | 1:50 |
| Interferon-γ | IFNγ | BV421 | XMG1.2 | BD | 563376 | 1:40 |
| Fc Block | CD16/CD32 | - | 2.4G2 | BD | 553141 | 1:50 |

### qRT-PCR and Western blotting

RNA was extracted, cDNA generated, and gene expression level determined by qRT-PCR as previously described in (70). Taqman Probes (Thermo 4331182): APOBEC3B, Hs00358981_m1; ACTB, Hs01060665_g1; Actb, Mm02619580_g1; Foxp3, Mm00475162_m1; Gzma, Mm01304452_m1; Tbx21, Mm00450960_m1; Prf1, Mm00812512_m1. mRNA expression was normalized to β-actin and presented as the relative fold change. To compare A3B expression levels between murine cell lines (SMF-A3B and EMT6-A3B) and human cell lines (BT474), A3B expression was not normalized to account for differences in β-actin expression between mouse and human cells; fold change of relative Ct value was presented.

For Western blotting, cells were treated doxycycline as described and harvested. Cells were lysed in RIPA buffer and 1x Halt Proteinase/Phosphatase Inhibitor (Invitrogen 78444). Protein concentration in the supernatant was determined by Bradford assay. Laemmli Sample Buffer (BioRad 1610747) was added to diluted protein samples and denatured at 95°C for 5 minutes. 20 µg of denatured protein was loaded into wells of 10-15% SDS-PAGE gel and ran at 90-125 V for 1 hour. Gel was transferred to immunoblot membrane using wet transfer at 90 V for 1 hour. Membranes were incubated with blocking buffer for 1 hour at room temperature and then primary antibodies at dilutions listed below overnight at 4°C. Membranes were washed in PBS-Tween 20 and incubated with secondary antibodies at dilutions listed below for 1 hour at room temperature in the dark. Membranes were then washed and imaged using a Li-Cor Odyssey infrared imaging system and analyzed in ImageStudio Lite software (Li-Cor Biosciences).

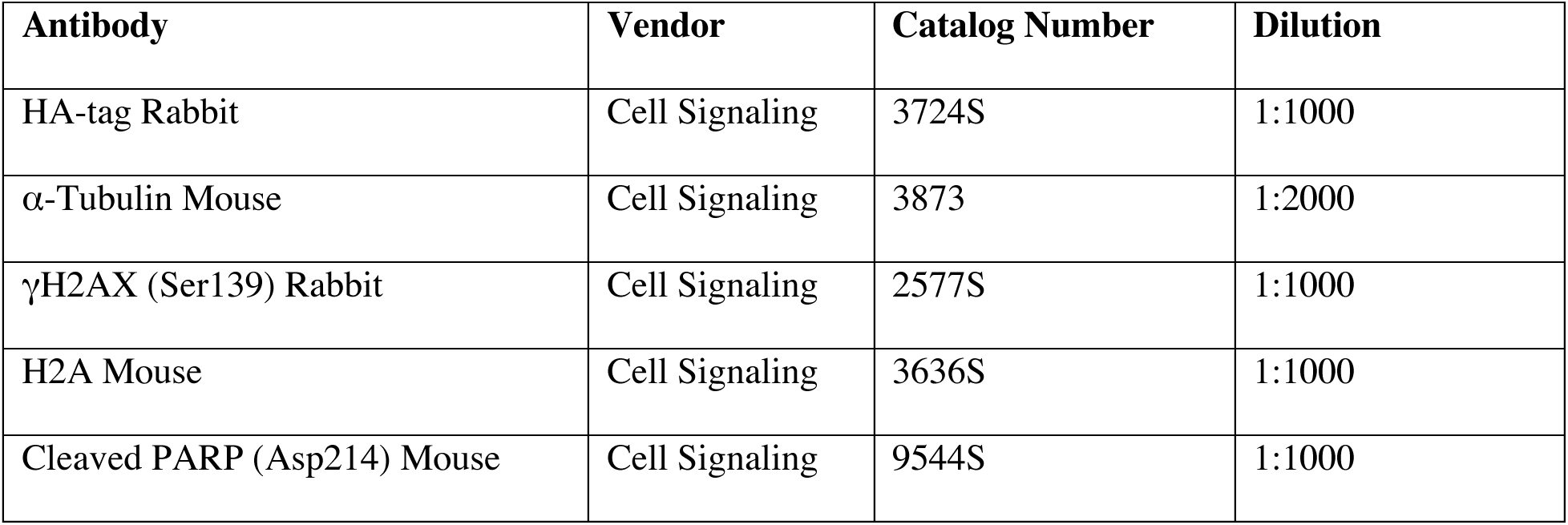

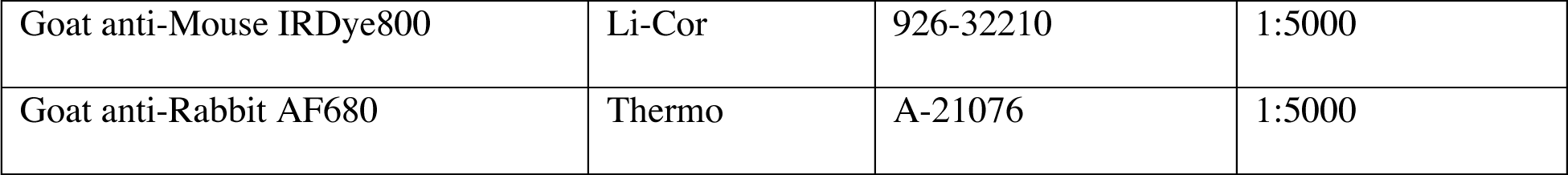

### Tissues, immunohistochemistry, and immunofluorescence

Tumors were harvested and fixed in 10% normal formalin overnight before paraffin-embedding for immunohistochemistry by Duke Pathology Research Immunohistology Lab (Duke University, Durham, NC). Slides were imaged at 4 fields of view per tumor with Zeiss Axio Imager Widefield fluorescence microscope.

Tumors were harvested and frozen in OCT for immunofluorescence staining. Slides were fixed in 4% paraformaldehyde for 10 minutes. Slides were washed in PBS, permeabilized in 0.5% Triton-X 100 for 20 minutes, washed in PBS, and blocked in 3% BSA and 10% normal goat serum for 1 hour at room temperature. Slides were incubated with primary antibodies listed below overnight at 4°C, washed, and incubated in with secondary antibodies listed below for 1 hour at room temperature. Slides were then washed in PBS, stained with DAPI for 10 minutes, and coverslips were mounted on slides with Prolong Gold (Thermo P36930). For γH2AX foci quantification, 8 fields of view were imaged per slide with Leica SP5 Inverted Confocal fluorescence microscope. For assessing expression of HA-tagged A3B in tumors, slides were imaged with Zeiss Axio Imager Widefield fluorescence microscope. Images were analyzed with ImageJ Fiji.

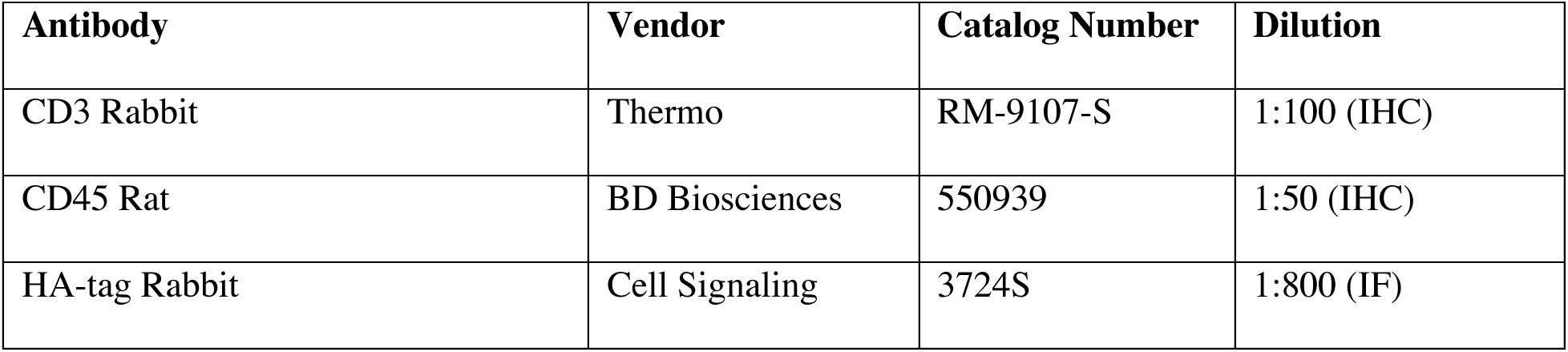

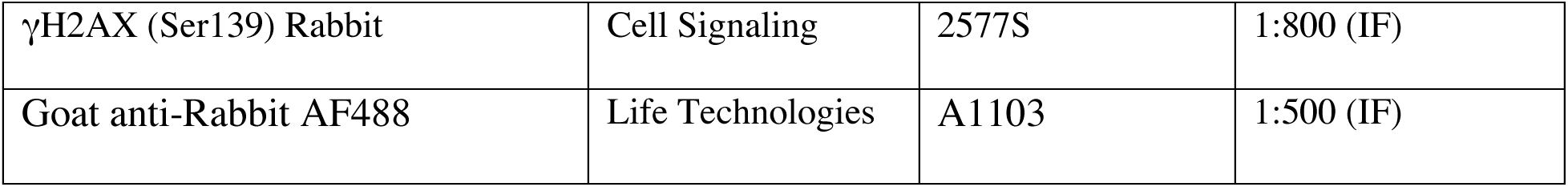

### Cytidine deaminase activity assay

Cells were treated with doxycycline as described and harvested. Cells were lysed for 10 minutes on ice in 25 mM HEPES (pH7.4, diluted in molecular grade water), 10% glycerol, 150 mM NaCl, 0.5% Triton X-100, 1 mM EDTA, 1 mM MgCl_2_, 1 mM ZnCl_2_, and 1:100 protease inhibitor (Sigma P8340). Protein concentration of the supernatant was determined using DC^TM^ Protein Assay (BioRad) and manufacturer protocol. 10 µg of protein was incubated for 2 hours at 37°C with 4 pmol of oligonucleotide listed below, 0.5 µL of uracil DNA glycosylase enzyme (NEB M0280S), 2 µL of 10x uracil DNA glycosylase buffer (NEB M0280S), 2.5 µL RNase A (M0280S) up to a 20 µL reaction volume with molecular grade H_2_0. Then 10 µL of 1N NaOH was added and heated to 95°C for 10 minutes to break the DNA backbone. Then 30 µL of 2x RNA loading dye was added and heated to 95°C for 3 minutes to denature the DNA. 15% Urea-TBE-PAGE gel was made with 3.75 mL of 40% Acryl (29:1), 4.8 g of ultra-pure urea, 1 mL of 10x TBE buffer, 5.25 mL of H_2_0, 99 µL of 10% APS, and 4 µL of TEMED. 15% Urea-TBE-PAGE gel was prewarmed for 1 hour at 150 V. 5 µL of denatured sample was added per well and ran at 150 V for 30-45 minutes. Gels were imaged with Li-Cor Odyssey infrared imaging system and analyzed in ImageStudio Lite software (Li-Cor Biosciences) to quantify the percent of deamination.

Oligonucleotide containing cytosine:

/5IRD700/ATTATTATTATTCAAATGGATTTATTTATTTATTTATTTATTT

Positive control oligonucleotide containing uracil:

/5IRD700/ATTATTATTATTUAAATGGATTTATTTATTTATTTATTTATTT

### RNA-sequencing and analysis

RNA was isolated from tumors using the RNeasy kit (Qiagen). RNA was sequenced using Stranded mRNA-seq libraries and the NovaSeq 6000 S1 sequencing platform with 50 bp paired-end reads by the Duke GCB Sequencing and Genomic Technologies Shared Resource (Duke University, Durham, NC).

RNA-seq data was trimmed with Trim Galore! (Galaxy Version 0.6.3; Krueger, F., Babraham Institute, http://www.bioinformatics.babraham.ac.uk/projects/trim_galore/) and then FastQC (Galaxy Version 0.72+galaxy1; Andrews, S. (n.d.). FastQC A Quality Control tool for High Throughput Sequence Data. Retrieved from http://www.bioinformatics.babraham.ac.uk/projects/fastqc/) was used to assess quality. Reads were aligned to the GRCm38 reference mouse genome using RNA STAR (Galaxy Version 2.7.5b)(71) and vM25 annotation file downloaded from the Gencode server. Reads were counted with featureCounts (Galaxy Version 1.6.4+galaxy2)(72) and differential gene expression analysis was performed with DESeq2 (Galaxy Version 2.11.40.6+galaxy1)(73).

Gene ontology (GO) analysis was performed using GO Ontology database (DOI: 10.5281/zenodo.4033054 Released 2020-09-10)(74, 75) for the log2 fold change of genes with a FDR adjusted p-value < 0.05. Gene set enrichment analysis (GSEA) was performed using GSEA v4.1.0(76, 77) using preranked gene list and Hallmark v7.2 gene set database at 1,000 permutations. *In silico* flow cytometry was used to compute immune cell fractions with CIBERSORTx(78) and the LM22 dataset.

### T cell receptor-sequencing and analysis

RNA was isolated from tumors using the RNeasy kit (Qiagen) and TCR beta chain libraries were generated using SMARTer Mouse TCR a/b Profiling Kit (Clontech). Samples were pooled to a final pool concentration of 4 nM and diluted to a final concentration of 13.5 pM, including a 5–10% PhiX Control v3 spike-in. Libraries were sequencing using MiSeq600 v3 300 bp paired-end reads. MiXCR(79) was used to calculate clonotype frequencies with recommended settings and vegan R package (Jari Oksanen, F. Guillaume Blanchet, Michael Friendly, Roeland Kindt, Pierre Legendre, Dan McGlinn, Peter R. Minchin, R. B. O’Hara, Gavin L. Simpson, Peter Solymos, M. Henry H. Stevens, Eduard Szoecs and Helene Wagner (2020). vegan: Community Ecology Package. R package version 2.5-7. https://CRAN.R-project.org/package=vegan) was used to calculate Shannon entropy diversity index. DE_50_ was calculated as the number of clonotypes occupying the top 50% of read counts, divided by the total number of read counts.

### ELISpot

Spleens and tumors were harvested from 4 APOBEC tumor-bearing mice. Single cell splenocytes suspensions were generated and cryopreserved. Tumor chunks were snap-frozen in liquid nitrogen. For co-culture, splenocytes were thawed at 37°C, washed 3 times in splenocyte medium (RPMI, 10% FBS, and 1% penicillin/streptomycin), and counted. Tumor chunks were thawed on ice, lysed using 4 rounds of −80°C freeze/thaw cycles, and protein concentrations were determined from the supernatant using Bradford assay. 1×10^6^ splenocytes from APOBEC tumor-bearing mice were co-cultured for 48 hours with 100 µg/mL of tumor lysate protein. 2.5×10^5^ splenocytes from naïve mice were co-cultured for 48 hours with 1 µg/mL of concanavalin A as a positive control or alone as a negative control. Mouse IFN-γ ELISpot PLUS kit (MABTECH 3321-4APT-2) manufacturer protocol was followed for development of the spots. Plates were dried overnight and imaged and quantified using CTL ImmunoSpot 7.0.26.0 software.

### Bioinformatics analysis of human breast cancers

To estimate immune cell infiltration, we quantified the gene expression of individual immune checkpoint genes or immune cell gene signatures (56, 57) collapsed into one value for each signature using a PCA-based method (see Supplementary Table 1 for gene lists). Raw gene counts from RNA-seq experiments for TCGA-BRCA patients were first queried from the National Cancer Institute Genomic Data Commons (GDC) (80) using the R package TCGAbiolinks (v2.12.6) (81), and normalized to effective library sizes calculated by the Trimmed Mean of M-values (TMM) (82) method and transformed by the voom method (83) implemented in the R packages edgeR (v3.28.0) (84) and Limma (v3.42.0) (85), respectively. For each gene signature, the first principal component (PC1) of a PCA model was used to summarize the gene expression values of the signature into a single score.

To calculate APOBEC enrichment score, single nucleotide polymorphisms (SNP) data called by the somatic mutation caller MuTect2 (86) were also queried from the GDC. For each tumor, an APOBEC mutagenesis enrichment score was calculated based on C>T mutations occurring in TCW motifs as described by Roberts et al (9).

Heatmaps of the relative expression of immune cell gene signatures in APOBEC-high and APOBEC-low tumors were created using R package Morpheus (https://software.broadinstitute.org/morpheus). Samples were grouped by subtype (HER2-enriched or basal-like), and Euclidian hierarchical clustering and cutting the dendrogram was used to identify 2 main immune clusters in the HER2-enriched subtype and 2 main immune clusters in the basal-like subtype (immune cluster 1 and cluster 2).

To measure the correlation of APOBEC enrichment score and immune gene signatures in samples based on subtype, Spearman’s rho was calculated, and the significance was determined by Spearman’s rank correlation test. P-values were adjusted for multiple testing using the Benjamini-Hochberg method to control the false discovery rate.

To assess differences in genetic heterogeneity between immune clusters, the number of subclonal mutations per TCGA sample was downloaded from (61).

### Gene expression-based classifier of APOBEC mutagenesis

APOBEC enrichment scores were calculated from TCGA tumors as in methods described above. We performed classification to nearest centroids to identify sets of genes that would distinguish individuals with high APOBEC enrichment from those without (87). After constructing a matrix of log-2 transformed, median-centered gene expression values for TCGA-BRCA samples, we filtered genes to the top 5% most differentially expressed (N = 1,026 genes) between APOBEC-high and APOBEC-low samples using the samr package (R. Tibshirani, Michael J. Seo, G. Chu, Balasubramanian Narasimhan and Jun Li (2018). samr: SAM: Significance Analysis of Microarrays. R package version 3.0. https://CRAN.R-project.org/package=samr). We performed 10-fold cross validation by randomly splitting the TCGA samples into 10 groups and training the classifier on nine of these groups (training set), leaving the remaining group to serve as an internal validation set (test set). In each of the 10 iterations of training, we varied the number of genes used to predict each APOBEC group from 1 to 50 and assessed model performance by calculating sensitivity and specificity in both training and test sets. Mean sensitivity compared to APOBEC enrichment calls derived from whole-exome-sequencing across each of the 10 folds ranged from 61-71% in training sets, with the maximum test set sensitivity reached at 5 genes per group (Supplementary Fig. S10A, B). We chose the final number of genes based on the maximum Youden’s index (sensitivity + specificity – 1). The maximum Youden’s index for test data was achieved using 5 genes per group (Y = 0.30), suggesting that a 10-gene classifier was optimal for prediction (Supplementary Fig. S10C). Applied to the full TCGA breast cancer cohort, the predictor achieved 69% sensitivity and 61% specificity against APOBEC enrichment calls from whole-exome sequencing data, for an overall accuracy of 63% (Supplementary Fig. S10D). This accuracy is consistent with what would be expected given the observed instability in signature detection when resampling mutations within an individual, particularly in contexts of low mutation frequency (41, 42). Finally, the expression-based predictor was applied to RNA-seq data from a sample of 12 mouse tumors from NSG mice (6 A3B-expressing tumors and 6 control tumors) to classify tumors demonstrating the APOBEC mutational signature. In this instance, sensitivity and specificity were calculated using A3B/control status as the gold standard.

### Statistical reporting

One-way ANOVA and Tukey’s multiple comparisons test was used to assess statistical significance of qRT-PCR gene expression, colony formation assay, and MHC-I expression by flow cytometry. One-way ANOVA and Sidak’s multiple comparisons test were used to assess statistical significance of the mouse tumor volume on a single day as indicated, differences in APOBEC enrichment score from human data, and the number of subclonal mutations in immune clusters from human data. Two-way ANOVA and Dunnett’s multiple comparisons test was used to determine the statistical significance of differential cell growth *in vitro* using CellTiter Glo assay. Two-way repeated-measures ANOVA and Tukey’s multiple comparison was used to measure statistical significance of changes in tumor volume over time *in vivo*. The adjusted p-values are reported for each.

Fisher’s exact test was used to assess differences in response to checkpoint inhibition (CR/PR by percent change in tumor volume from treatment start day). Student’s t-test was used to test the statistical significance of differences in tumor mass at endpoint, flow cytometry, IHC/IF, and TCR-seq diversity measurements. Student’s t-test p-values are reported. Statistical analysis was performed and graphs were created in R version 4.0.2 or using GraphPad Prism version 8.0.1.

### Research reproducibility

Source code to reproduce analyses of gene expressed-based APOBEC classifier and to reproduce analyses of APOBEC enrichment and quantification of immune signature gene expression is available at https://github.com/ashleydimarco/alvarezlab-APOBEC

## Author contributions

**Ashley V. DiMarco:** conceptualization, data curation, formal analysis, supervision, validation, investigation, visualization, methodology, writing—original draft, project administration; **Xiaodi Qin:** data curation, formal analysis, methodology, writing—review and editing for APOBEC enrichment score, immune gene signatures, and correlation analysis of TCGA data; **Sarah Van Alsten:** data curation, formal analysis, methodology, writing—review and editing for gene expressed-based classifier of APOBEC mutagenesis; **Brock McKinney:** data curation, resources; **Nina Marie G. Garcia:** data curation, writing—review and editing; **Jeremy Force:** methodology; **Brent A. Hanks:** supervision, writing—review and editing for depletion and checkpoint inhibitor studies; **Melissa A. Troester:** supervision, writing—review and editing for gene expressed-based classifier of APOBEC mutagenesis; **Kouros Owzar:** supervision, writing—review and editing for APOBEC enrichment score, immune gene signatures, and correlation analysis of TCGA data; **Jichun Xie:** supervision, writing—review and editing for APOBEC enrichment score, immune gene signatures, and correlation analysis of TCGA data; **James V. Alvarez:** conceptualization, supervision, funding acquisition, project administration, writing—review and editing.

## Acknowledgements

We thank Dr. Lewis Chodosh (University of Pennsylvania) for providing the SMF cell line. We thank Dr. Michael Plebanek and Dr. Nicolas Devito (Duke University) for technical advice with flow cytometry and ELISpot. We thank Elizabeth Mendes and Alexandra Bennion (Duke University) for providing technical assistance, and Dr. Andrea Walens (University of North Carolina at Chapel Hill) for reviewing the manuscript. We thank the Duke Pathology Research Immunohistology Lab for paraffin processing and IHC staining of tissue. We thank Dr. Nicolas Devos (Duke University) and the Duke University School of Medicine Sequencing and Genomic Technologies Shared Resource for providing library preparation and sequencing for RNA-seq and WES analyses, and sequencing for TCR-seq analysis. This work was funded by the National Cancer Institute under award R01CA208042 (to J.V.A.) and T32-CA009111 (to A.V.D.), as well as the American Cancer Society under award 132556-RSG-18-130-CCG (to J.V.A.) and by startup funds from the Duke Cancer Institute, the Duke University School of Medicine, the Whitehead Foundation (to J.V.A.), and the National Institutes of Health under T32-GM007184 (to A.V.D.).

## SUPPLEMENTARY MATERIAL

**Supplementary Figures S1-S10**

**Supplementary Table S1**: Spreadsheet of gene lists for immune cell gene signatures for analyses in Figure 6 and Supplementary Figure S9. The genes that were absent from the TCGA-BRCA RNA-seq dataset are colored in red.

**Supplementary Table S2**: Spreadsheet containing TCGA-BRCA patient ID and data used for analyses in Figure 6 and Supplementary Figure S9. Column descriptions:

Sample_ID – TCGA sample identifier

Cluster_Number – APOBEC-high or -low immune cluster number (e.g. “APOBEC-high HER2-1” refers to APOBEC-high HER2 subtype Immune Cluster 1)

Age_Median – patient age

ER.Status – clinical ER status

PR.Status – clinical PR status

Her2.Status – clinical

HER2 status PAM50 – PAM50 subtype

Pathologic_stage – clinical pathologic stage

Histological_type – clinical histological type

n_C_mut – number of C>T/G (or G>A) mutations

n_C_con – number of C (or G) within the 41-nucleotide region centered on the C>T/G (or G>C/A) mutations

n_TCW_mut – number of C>T/G (or G>C/A) mutations in TCW (or WGA) motifs

n_TCW_con – number of TCW (or WGA) motifs within the 41-nucleotide region centered on the mutated motifs, TCW to TTW/TGW (or WGA to WAA/WCA).

APOBEC – APOBEC enrichment score

Number_of_Subclonal_Mutations – number of subclonal mutations from Raynaud et al. 2018 The remaining columns are principal component analysis (PCA)-collapsed log2 normalized gene expression of immune cell gene signatures from RNA-seq data.

**Supplementary Table S3**: Spreadsheet of correlations between APOBEC enrichment score and immune cell signatures used for analyses in Figure 6. Column descriptions:

PAM50 – PAM50 subtype

rho – Spearman’s rho value from correlation analysis

pvalue – p-value from correlation analysis

adjusted_pvalue – adjusted p-value from correlation analysis

---

**Supplementary Figure S1:**
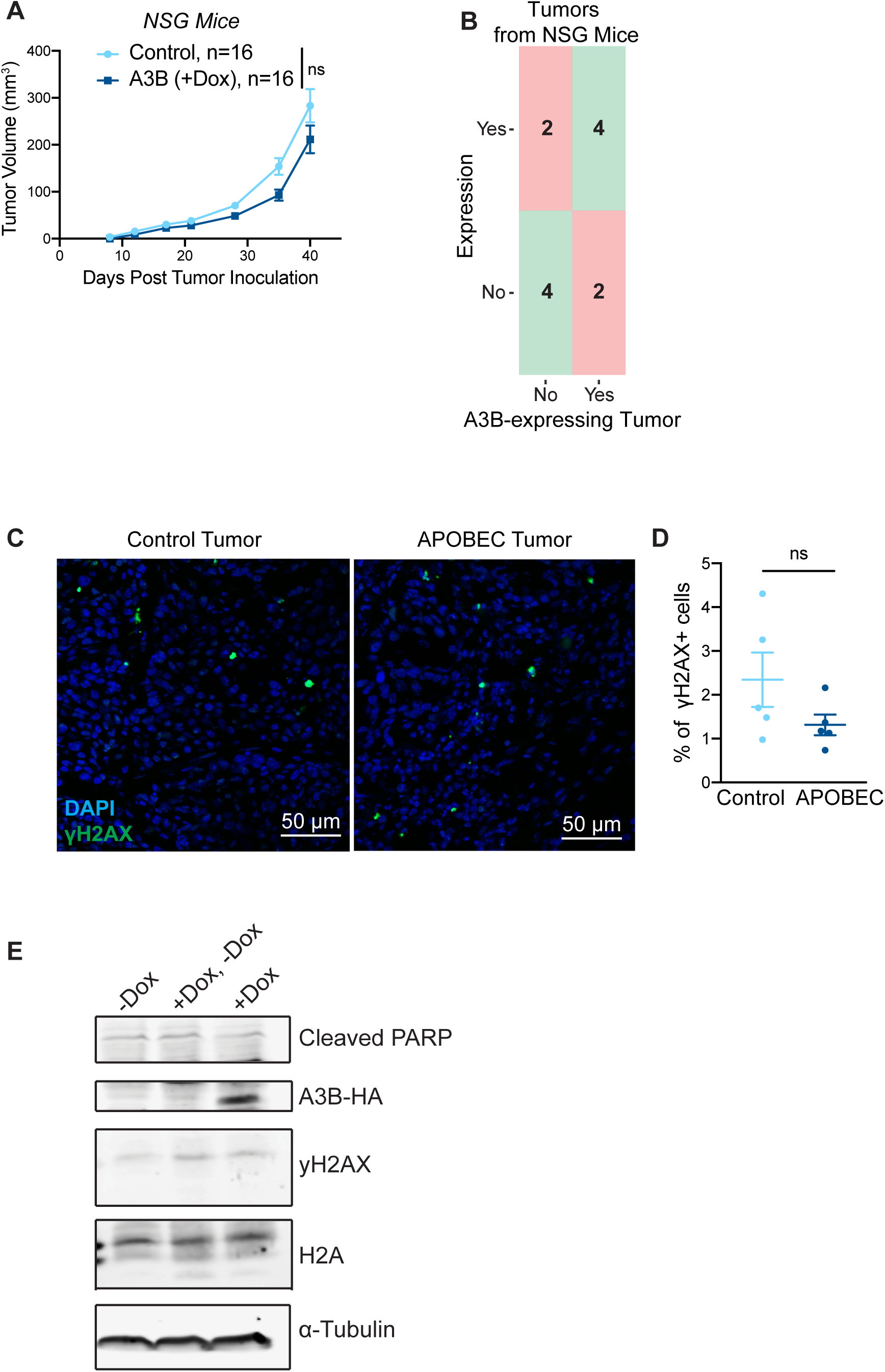
A3B expression does not alter tumor growth in immunodeficient mice and induces an APOBEC mutational gene expression signature without activating the DNA damage response. **(A)** Tumor volume (mm3) over time for control (n=16) and A3B-expressing tumors (+dox in drinking water; n=16) growing in NSG mice. Error bars denote mean ± SEM. Statistical significance was determined by two-way repeated-measures ANOVA and Tukey’s multiple comparisons test with the same control cohort as in Supplementary Fig. S5J. **(B)** Confusion matrix of 10 gene predictor applied to sample of 12 mouse tumors in NSG mice (6 A3B-expressing tumors and 6 control tumors). Squares in red (upper left and bottom right) denote incorrect classifications and squares in green (upper right and bottom left) represent correct classifications. Sensitivity and specificity were both 66%. **(C-D)** Immunofluorescence staining for γH2AX on control and APOBEC tumors growing in wildtype mice (see Figure 2B). Representative images are shown in (C) and quantification of γH2AX+ foci (number of foci/number of cells per field of view) is shown in (D). Five tumors per cohort were analyzed and 8 fields of view were averaged per tumor. DAPI is in blue and γH2AX is in green. Error bars denote mean ± SEM and statistical significance was determined by unpaired Student’s t-test. **(E)** Western blot analysis of HA-epitope tagged A3B, γH2AX, and cleaved PARP in SMF-A3B cells treated with or without dox for 2 weeks. α-Tubulin and histone H2A are shown as loading controls. ns > 0.05

**Supplementary Figure S2:**
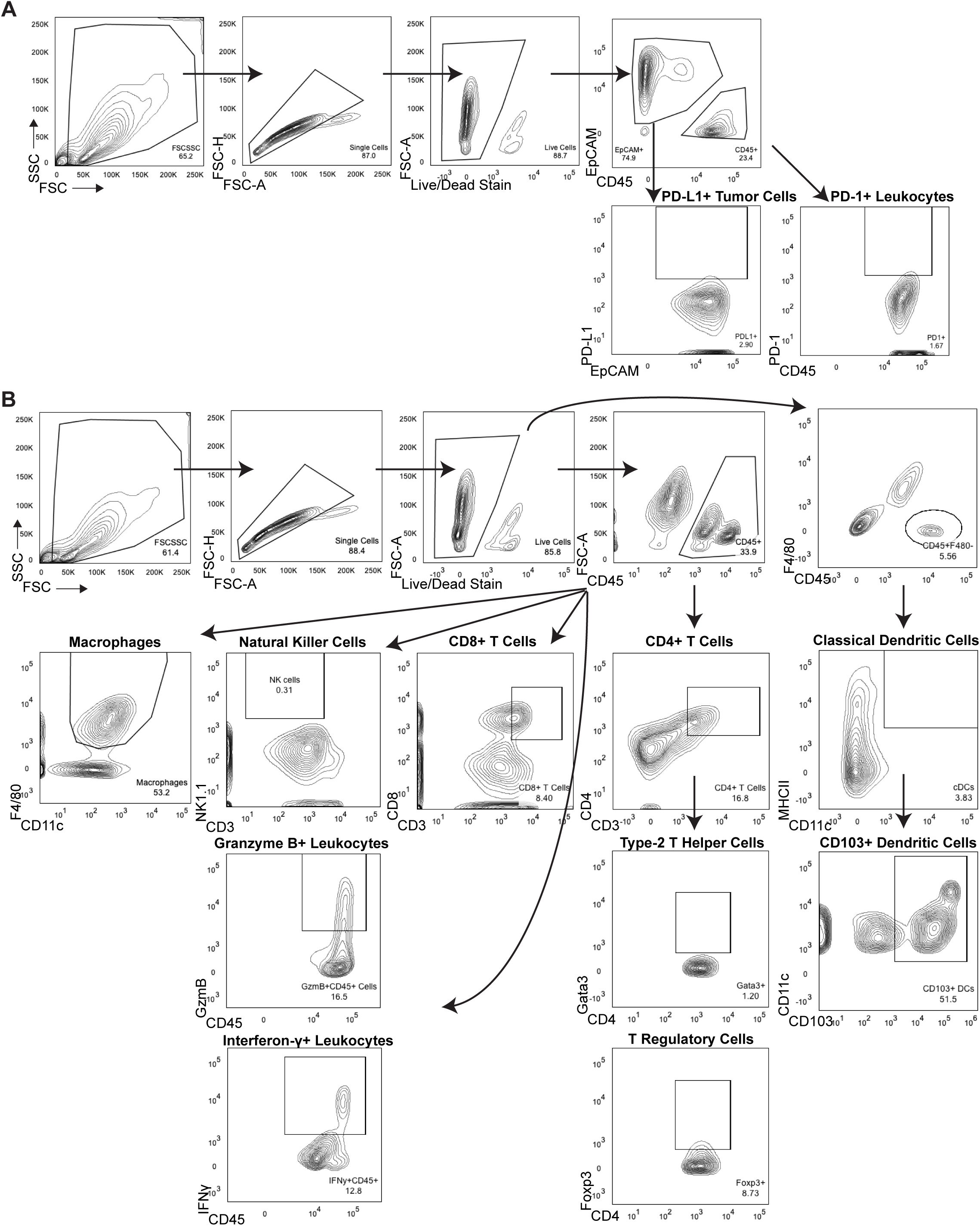
Flow cytometry gating strategy. **(A)** Gating strategy for PD-1 and PD-L1 expression on immune cells and tumor cells. **(B)** Gating strategy for macrophages, natural killer cells, granzyme B+ immune cells, interferon-γ+ immune cells, CD8+ T cells, CD4+ T cells, type-2 T helper cells, T regulatory cells, and CD103+ dendritic cells.

**Supplementary Figure S3:**
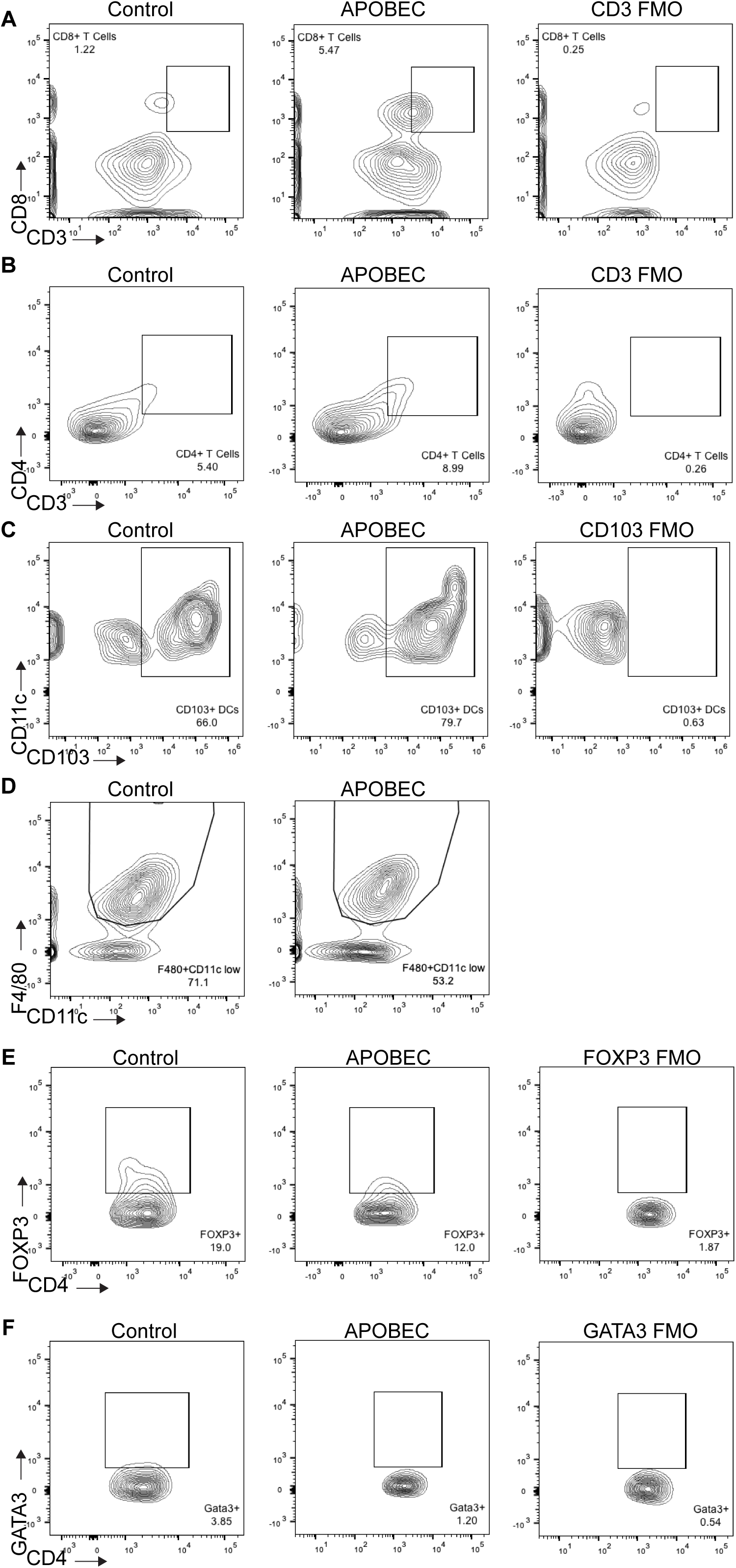
Representative FACS plots showing immune cell infiltration in APOBEC tumors. **(A)** Staining for CD8+ T cells (CD8+CD3+) and CD3 fluorescence minus one (FMO) control without CD3 antibody. **(B)** Staining for CD4+ T cells (CD4+CD3+) and CD3 FMO. **(C)** Staining for CD103 expression on dendritic cells (CD103+CD11c+) and CD103 FMO. **(D)** Staining for macrophages (F4/80 +CD11clow). **(E)** Staining for T regulatory cells (CD4+FOXP3+) and FOXP3 FMO. **(F)** Staining for Type-2 T helper cells (CD4+GATA3+) and GATA3 FMO.

**Supplementary Figure S4:**
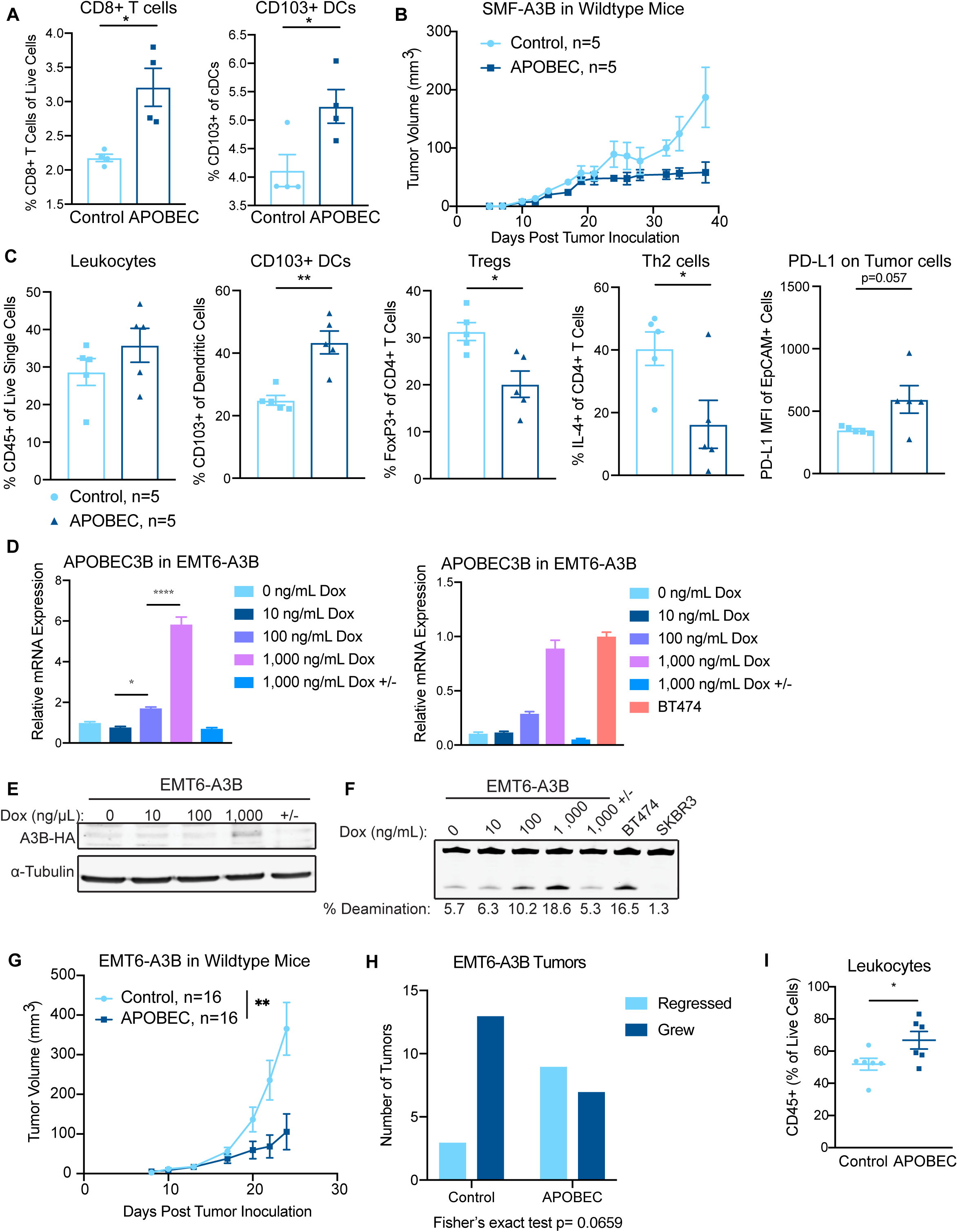
Tumor growth inhibition and increased immune infiltration in APOBEC tumors is reproducible and generalizable. **(A)** Bilateral tumor-draining inguinal lymph nodes (TDLN) were harvested from mice in Figure 2B and aggregated from 4 mice per cohort for flow cytometry. APOBEC TDLNs had increased CD8 + T cells and CD103+ DCs compared to control TDLNs. Error bars denote mean ± SEM and statistical significance was determined by unpaired Student’s t-test. **(B)** Tumor volume (mm3) over time for unilateral control tumors (n=5), and APOBEC tumors (n=5) generated from SMF-A3B cells in wildtype mice in an independent experiment, demonstrating that the growth defect of APOBEC tumors is reproducible in an independent experiment. Error bars denote mean ± SEM. **(C)** Control tumors (n=5) and APOBEC tumors (n=5) from (B) were harvested and immune profiled by flow cytometry. Quantification shows the APOBEC tumors had increased leukocytes, CD103+ dendritic cells (DCs), and tumor cell PD-L1 expression (MFI, mean fluorescence intensity), while T regulatory cells (Tregs) and type-2 T helper (Th2) cells were reduced in APOBEC tumors. These results demonstrate that immune infiltration in APOBEC tumors is reproducible in an independent experiment. Error bars denote mean ± SEM and statistical significance was determined by unpaired Student’s t-test. **(D)** qRT-PCR analysis for APOBEC3B expression in EMT6-A3B cultured with or without dox for 2 days. 1,000 ng/mL Dox +/− indicates cells cultured with 1,000 ng/mL dox for 2 days followed by removal of dox for 3 days prior to analysis. Left: A3B expression relative to 0 µg/mL dox condition. Right: A3B expression relative to BT474 cells. Results show 3 biological replicates and error bars depict mean ± SEM. Significance was determined using a one-way ANOVA and Tukey’s multiple comparisons test. **(E)** EMT6-A3B cells were cultured as in (D) and cell lysates were harvested for western blot of HA-tagged A3B protein. **(F)** EMT6-A3B cells were cultured as in (D) and cell lysates harvested for in vitro deaminase activity assay. Deaminase activity is comparable to that of human cell line, BT474. SKBR3 human cell line is A3B-null and shown as a negative control. **(G)** Tumor volume curves for control (-dox; n=16) and APOBEC (+dox; n=16) tumors derived from EMT-A3B cells orthotopically implanted in the mammary gland of syngeneic BALB/c mice. Error bars denote mean ± SEM and statistical significance was determined by two-way repeated-measures ANOVA. **(H)** The fraction of control and APOBEC EMT6 tumors that grew or spontaneously regressed following tumor cell injection. Fisher’s exact test, p=0.0659. **(I)** Flow cytometry quantification of leukocytes in control (n=6) and APOBEC (n=6) EMT6 tumors from (G). Error bars denote mean ± SEM and statistical significance was determined by unpaired Student’s t-test. * p < 0.05, ** p < 0.01, **** p < 0.0001

**Supplementary Figure S5:**
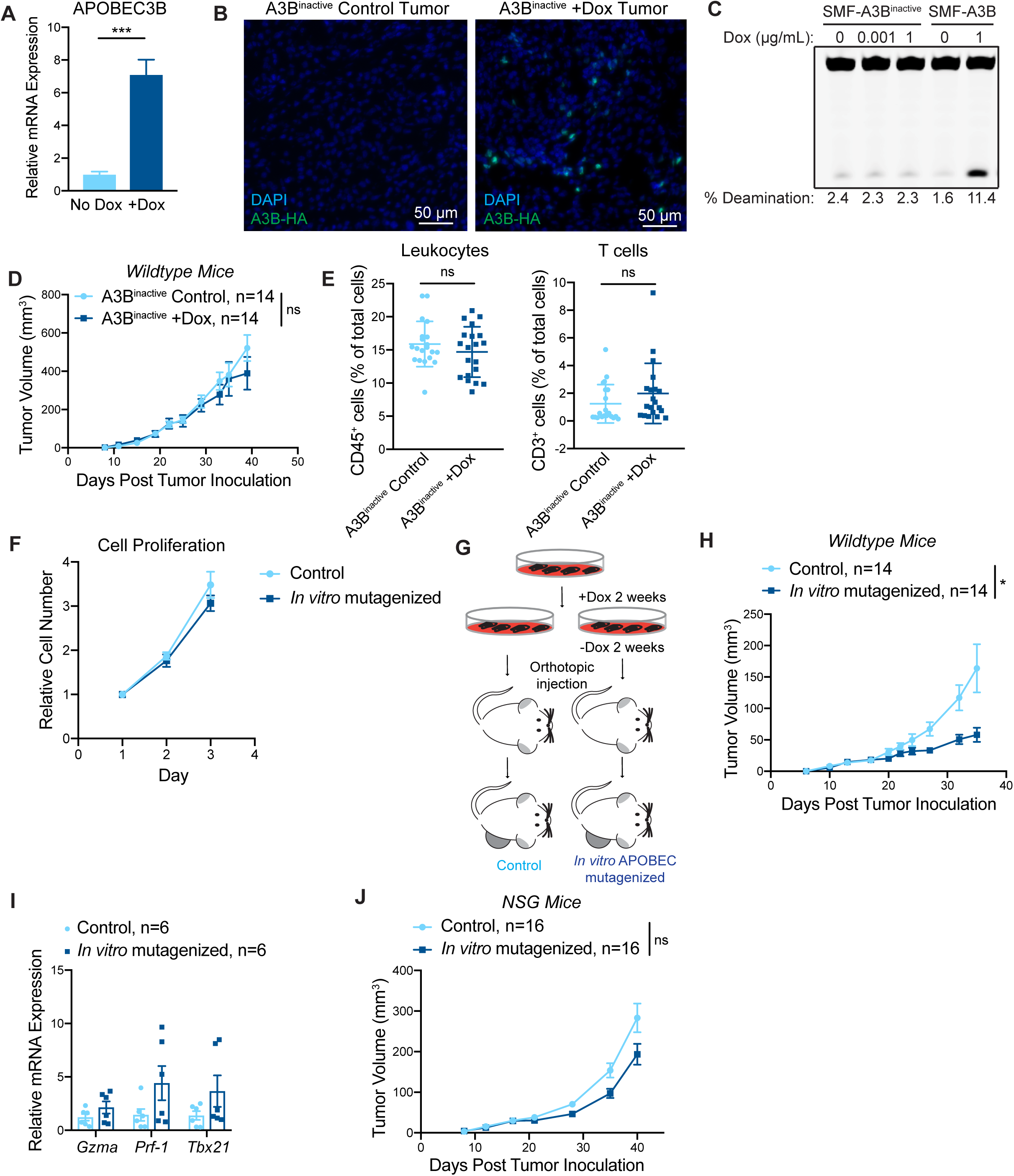
The APOBEC tumor growth defect requires the catalytic activity of A3B. **(A)** qRT-PCR of A3B gene expression in SMF-A3B^inactive^ cells treated with 1 ug/mL dox for 5 days. Error bars denote mean ± SD for 3 technical replicates and statistical significance was determined by unpaired Student’s t-test. **(B)** Immunofluorescence staining for HA epitope-tagged A3B in control tumors (-dox) and tumors expressing A3B^inactive^ (+dox). **(C)** In vitro deaminase activity assay in SMF-A3B^inactive^ cells treated with dox. SMF-A3B cells are shown as a control. **(D)** Tumor volume (mm3) over time for control tumors (-dox; n=14) and tumors expressing A3B^inactive^ (+dox ;n=14) in wildtype mice. Error bars denote mean ± SEM. Statistical significance was determined by two-way repeated-measures ANOVA. **(E)** Quantification of IHC staining for CD45 (left) or CD3 (right) in control tumors (n=5) and tumors expressing A3B^inactive^ (n=5). Four fields of view were quantified for each tumor. Error bars denote mean ± SD. Statistical significance was determined by unpaired Student’s t-test. **(F)** Growth curves for control and in vitro APOBEC mutagenized cells. Data are shown as mean ± SD of 4 replicates. **(G)** Schematic showing experimental design for tumor growth experiment. SMF-A3B cells were cultured with or without dox for 2 weeks, then dox was removed for 2 weeks. These in vitro APOBEC mutagenized cells or control cells were orthotopically implanted in the mammary gland of mice in the absence of dox. **(H)** Tumor volume (mm3) over time for control (n=14) and in vitro APOBEC mutagenized tumors (n=14) in wildtype mice. Error bars denote mean ± SEM. Statistical significance was determined by two-way repeated-measures ANOVA. **(I)** qRT-PCR analysis for Granzyme A (Gzma), Perforin-1 (Prf-1), and T-bet (Tbx21) in control (n=6) and in vitro APOBEC mutagenized tumors (n=6). All genes showed a trend toward increased expression in the in vitro APOBEC mutagenized cohort that did not reach statistical significance. **(J)** Tumor volume (mm3) over time for control (n=16) and in vitro APOBEC mutagenized tumors (n=16) in NSG mice. Error bars denote mean ± SEM. Statistical significance was determined by two-way repeated-measures ANOVA and Tukey’s multiple comparisons test. Note that control mice are the same as in S1A. ns p > 0.05, ** p < 0.01, *** p < 0.001

**Supplementary Figure S6:**
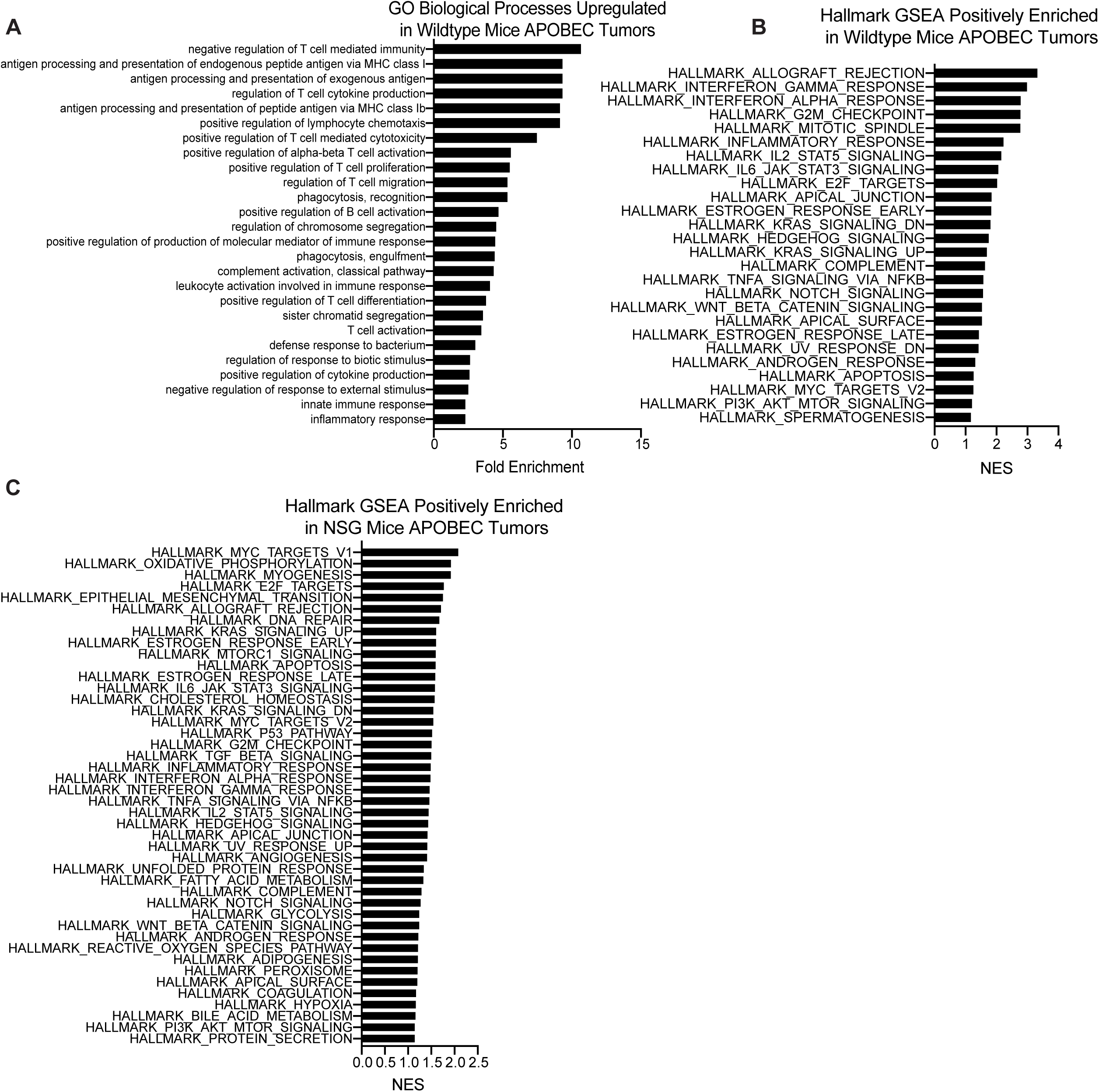
Gene expression analysis of control and APOBEC tumors. **(A)** All statistically significant main GO biological processes upregulated in APOBEC tumors from wildtype mice (FDR adjusted p < 0.05 by Fisher’s test). Fold enrichment is shown for each. **(B)** All statistically significant GSEA Hallmark pathways positively enriched in APOBEC tumors from wildtype mice (FDR adjusted p < 0.25). Normalized enrichment score (NES) is shown for each. **(C)** All statistically significant GSEA Hallmark pathways positively enriched in APOBEC tumors from NSG mice (FDR adjusted p < 0.25). Normalized enrichment score (NES) is shown for each.

**Supplementary Figure S7:**
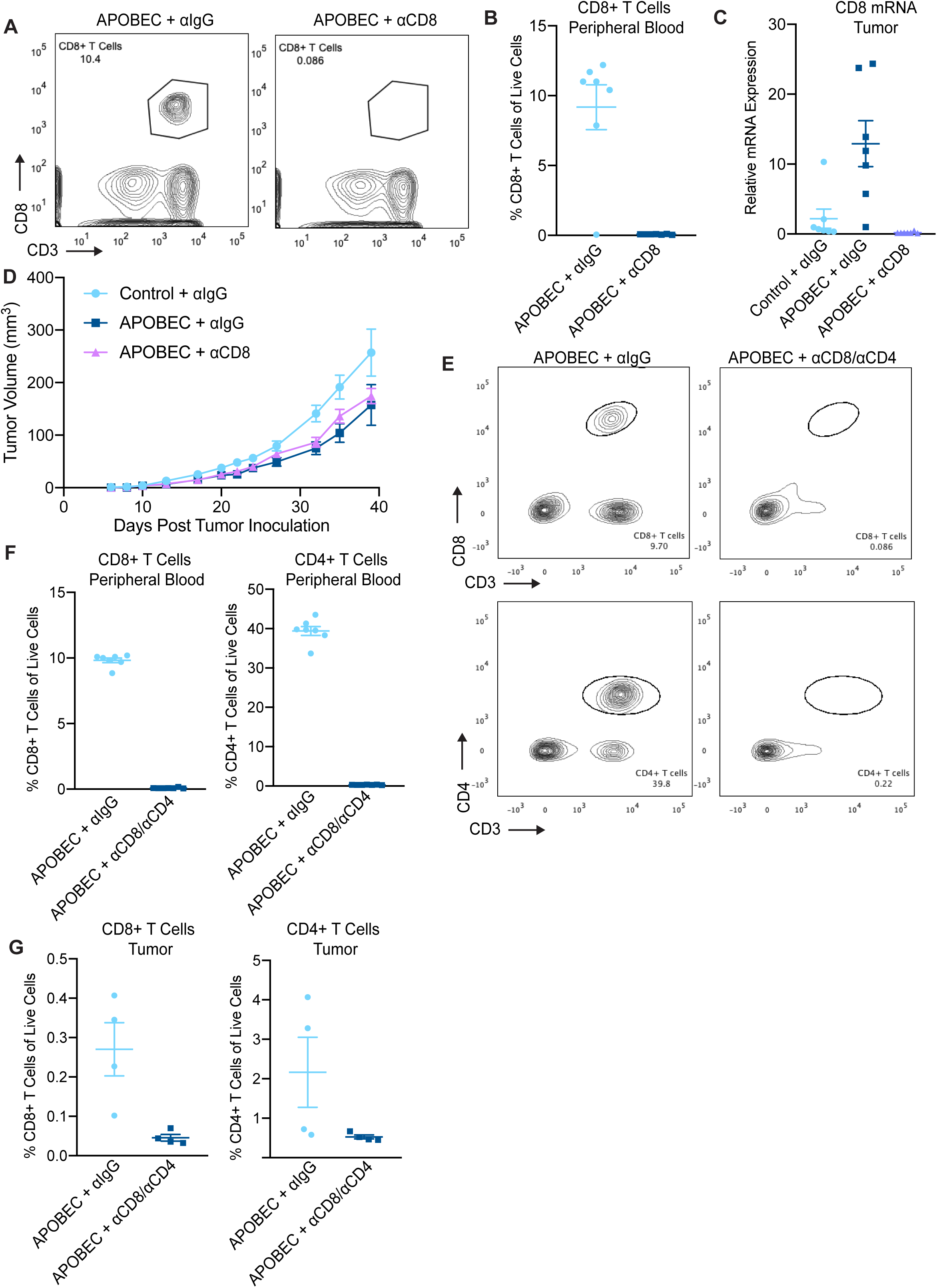
CD4+/CD8+ T cell depletion but not CD8+ T cell depletion alone rescues the growth defect of APOBEC tumors. **(A)** Flow cytometry showing depletion of CD8+ T cells in peripheral blood following intraperitoneal injection of an in vivo CD8 depleting antibody (300 µg/dose) or isotype-control antibody twice weekly. Peripheral blood was assayed on day 9 post tumor inoculation. **(B)** Quantification of CD8+ T cells in peripheral blood of isotype-control antibody treated mice (n=7) and αCD8 antibody treated mice (n=7) as in (A). Error bars denote mean ± SEM. **(C)** qRT-PCR for CD8 expression in tumors from the indicated cohorts: control tumors + αIgG (n=7); APOBEC tumors + αIgG (n=7); APOBEC tumors + αCD8 (n=7). Error bars denote mean ± SEM. **(D)** Tumor volume (mm3) over time for control + αIgG (n=14), APOBEC + αIgG (n=14), and APOBEC + αCD8 (n=14) tumors in wildtype mice. Error bars denote mean ± SEM. **(E)** Flow cytometry showing depletion of CD8+ and CD4+ T cells in peripheral blood following intraperitoneal injection of CD8 and CD4 depleting antibodies (200 µg CD8 and 200 µg CD4/dose) or isotype-control antibody twice weekly. Peripheral blood was assayed on day 25 post tumor inoculation. **(F)** Quantification of CD4+ and CD8+ T cells in peripheral blood of isotype-control antibody treated mice (n=7) and αCD8/αCD4 antibody treated mice (n=7) as in (E). Error bars denote mean ± SEM. **(G)** Flow cytometry quantification of CD8+ and CD4+ T cells in APOBEC tumors treated with isotype-control antibody or αCD8/αCD4 depleting antibodies. Error bars denote mean ± SEM.

**Supplementary Figure S8:**
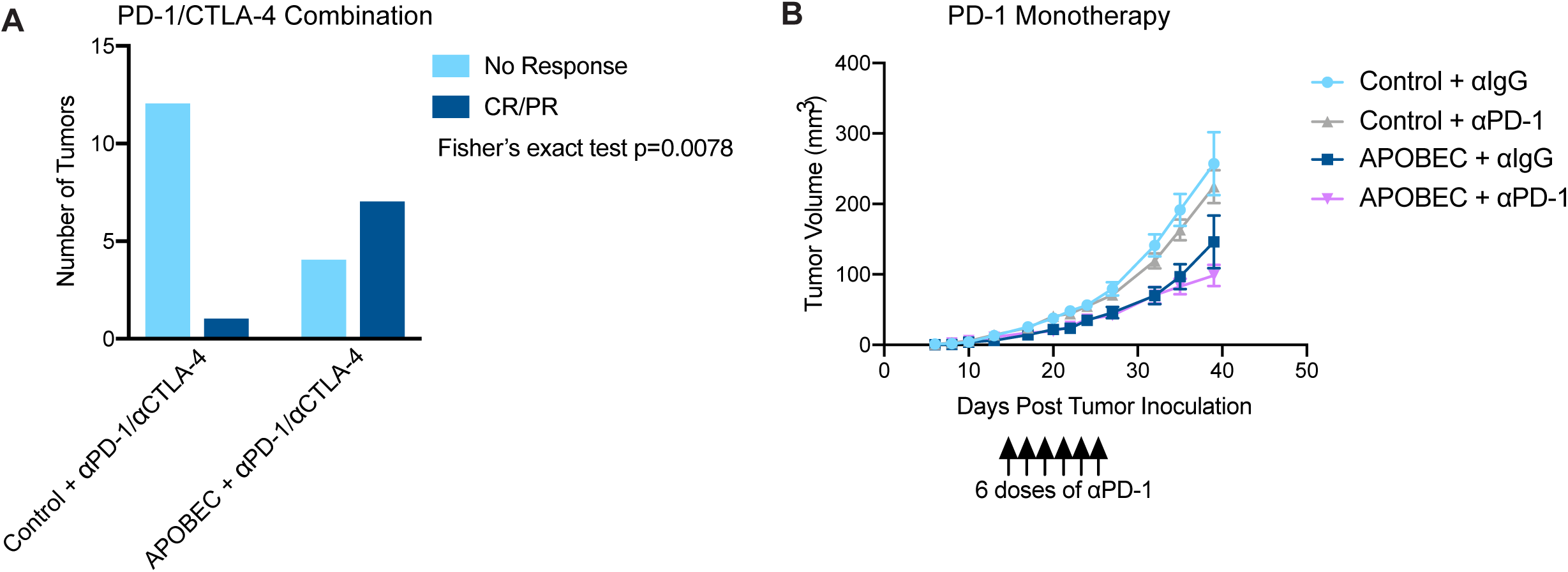
APOBEC activity renders HER2-driven mammary tumors responsive to combination anti-PD-1/anti-CTLA-4 therapy, but not anti-PD-1 monotherapy. **(A)** Response of control and APOBEC tumors to combination PD-1/CTLA-4 therapy. Number of tumors with no response or complete response/ partial response (CR/PR) are depicted. Statistical significance was determined by Fisher’s exact test (p=0.0078). **(B)** Tumor volume (mm3) over time for Control + αIgG (n=14), Control + αPD-1 (n=14), APOBEC + αIgG (n=14), and APOBEC + αPD-1 (n=14) tumors in wildtype mice. Mice were administered 6 doses of 200 µg of PD-1 or IgG isotype antibody on day 13, 15, 17, 20, 22, 24 post-tumor inoculation. Error bars denote mean ± SEM.

**Supplementary Figure S9:**
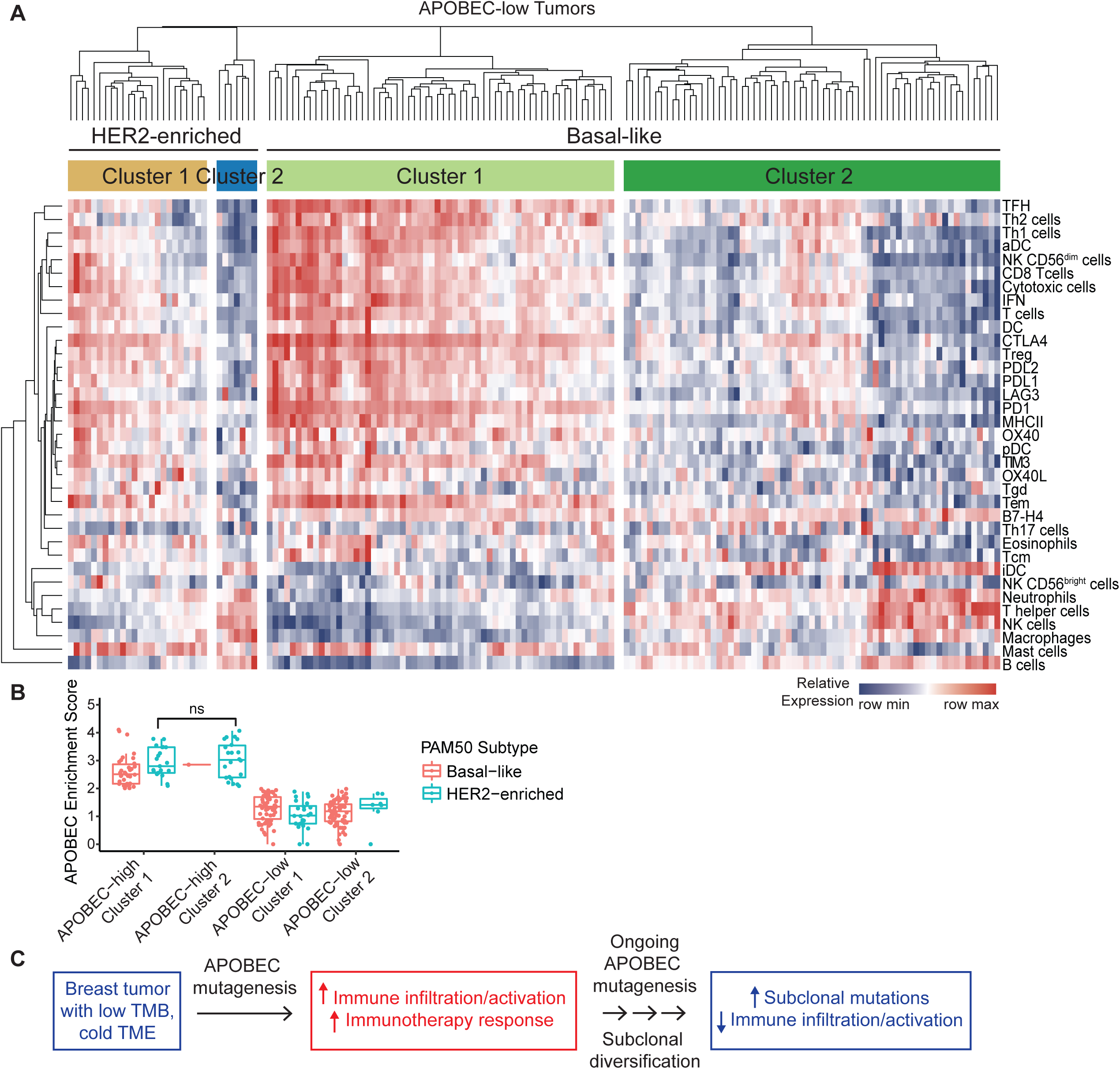
APOBEC-low TCGA tumors and immune signature clusters. **(A)** Heatmap showing the relative expression of immune cell gene signatures from TCGA RNA-seq data in APOBEC-low tumors, grouped by breast cancer subtype. Columns are individual patient tumors and rows are different immune cell gene signatures. Legend shows colors corresponding to relative expression levels (red, row max; blue, row min). Hierarchical clustering segregated tumors into 2 main clusters in the HER2-enriched subtype and 2 clusters in the basal-like subtype. **(B)** APOBEC enrichment score plotted for Basal-like and HER2-enriched tumors from each cluster. Boxplots show 25th percentile, median, and 75th percentile, while whiskers show minimum to maximum values excluding outliers. Statistical significance was determined by one-way ANOVA and Sidak’s multiple comparisons test. **(C)** Schematic of a model showing APOBEC mutagenesis increases immune activation, infiltration, and immunotherapy response in mouse and human breast tumors. But ongoing APOBEC mutagenesis can also generate subclonal diversification, which leads to increased subclonal mutations and decreased immune activation. ns p > 0.05

**Supplementary Figure S10:**
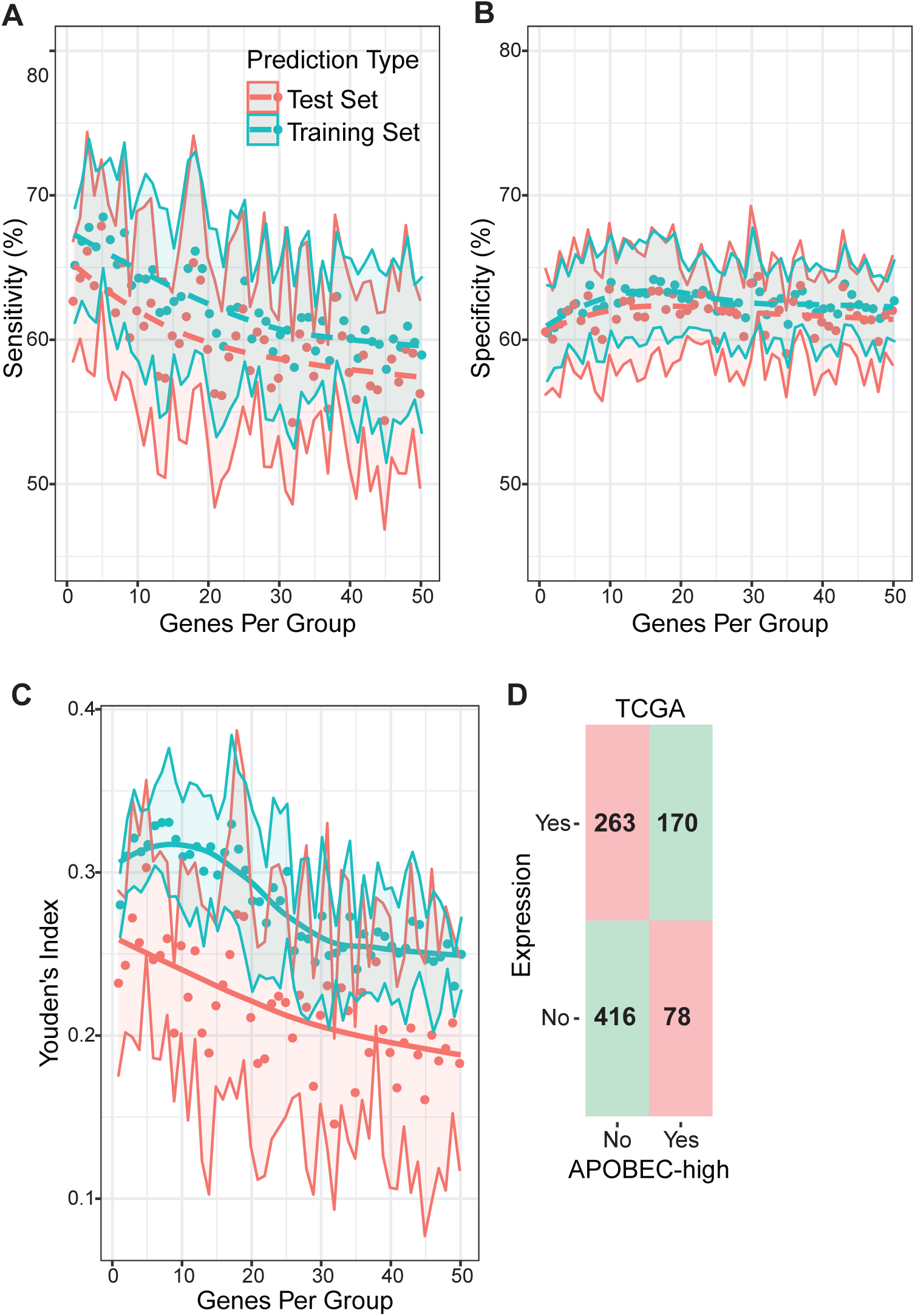
Methods for gene expressed-based classifier of APOBEC mutational signatures. **(A-B)** Ten-fold cross validated sensitivity (A) and specificity (B) of TCGA classification to nearest centroids (ClaNC) predictor using 1 to 50 of the top 5% most variably expressed genes to predict high versus low APOBEC enrichment by WES. In each run of the cross validation, 90% of all tumors were randomly selected to serve as the training set and the remaining 10% served as the test set. Points represent the mean sensitivity and specificity across the 10 folds, and confidence bands show standard deviation. **(C)** Ten-fold cross validated Youden’s index (sensitivity + specificity - 1) for test and training sets. The maximum Youden’s index in the test set was reached using 5 genes per group (10 total) and was therefore selected for the final model. Points represent mean Youden’s index, and confidence bands show standard deviation. **(D)** Confusion matrix of predicted (gene expression classifier) versus true (APOBEC enrichment score > 2 by WES) classifications in the full TCGA dataset using the 10 gene predictor. Squares in red (upper left and bottom right) denote incorrect classifications and squares in green (upper right and bottom left) represent correct classifications.

